# Distinct causes of three phenotypic hallmarks of hematopoietic aging

**DOI:** 10.1101/2025.01.14.632900

**Authors:** Jennifer J. Chia, Apeksha Singh, Yu-Sheng (Eason) Lin, Noa Popko, David Mastro, Yi Liu, Tiffany Tran, Jennifer K. King, Dinesh S. Rao, Alexander Hoffmann

## Abstract

Hematopoietic aging is characterized by chronic inflammation associated with myeloid bias, HSC accumulation, and functional HSC impairment. Yet it remains unclear how inflammation promotes these aging phenotypes. NFκB both responds to and directs inflammation, and we present an experimental model of elevated NFκB activity (“IκB^−^”) to dissect its role in hematopoietic aging phenotypes. We found that while elevated NFκB activity is not sufficient for HSC accumulation, HSC-autonomous NFκB activity impairs their functionality, leading to reduced bone marrow reconstitution. In contrast, myeloid bias is driven by the IκB^−^ proinflammatory bone marrow milieu as observed functionally, epigenomically, and transcriptomically. A new scRNA-seq HSPC labeling framework enabled comparisons with aged murine and human HSC datasets, documenting an association between HSC-intrinsic NFκB activity and quiescence, but not myeloid bias. These findings delineate separate regulatory mechanisms that underlie the three hallmarks of hematopoietic aging, suggesting that they are specifically and independently therapeutically targetable.

**Summary:** Aging is associated with inflammation and three hallmark hematopoietic dysfunctions. Using mouse models and single-cell analyses we report that each has a different cause: HSC accumulation is NFκB-independent, HSC functional impairment is driven by NFκB within HSCs, and myeloid bias is driven by the NFκB-altered milieu.

## INTRODUCTION

Phenotypes of hematopoietic aging include chronic inflammation, bias of hematopoietic output toward myeloid lineages and away from lymphoid fates, and hematopoietic stem cell (HSC) dysfunction.^1–3^ Aged HSCs accumulate in number, yet display increased quiescence and impaired responses to regenerative cues.^4–6^ Furthermore, HSCs and their multipotent progenitor (MPP) progeny (together HSPCs), display myeloid bias during diverse physiologic and pathologic inflammatory responses.^7–11^ While chronic inflammation is tightly linked with hematopoietic aging, whether it is a driver or consequence of myeloid bias, HSC functional decline, or HSC accumulation remains unclear.

NFκB is a ubiquitous regulator of inflammation, but is also downstream of inflammatory stimuli with roles in development, stress-responses, cancer. Interestingly, there is increased NFκB activity in aged HSCs.^12–20^ The NFκB signaling system, composed of IKK signal integration complexes, IκB negative regulators, and NFκB transcription factor (TF) dimers, is activated by tissue damage and inflammatory molecules (e.g. DAMPs, PAMPs, cytokines), culminating in NFκB nuclear translocation and gene transcription.^12,21^ Key transcriptional targets include cytokines (e.g. *Tnf*, *Il1a*), chemokines (e.g. *Ccl5*), and growth factors (e.g. *Csf3*), in addition to cell cycle, apoptosis and cell stress pathway components.^12,14,22–25^ Furthermore, the NFκB system also participates in the control of other signaling pathways such as MAP kinase, PI3 kinase, TGFβ, Wnt and type I interferon that also impact hematopoiesis.^22^ Both HSCs and their niches are altered in hematopoietic aging,^2,7,9,26,27^ yet there remains controversy regarding the contributions of each compartment to aging phenotypes.^28^ The diverse functions of NFκB across tissue types raises the question of whether NFκB-mediated signaling within both HSCs and their niches may each contribute to hematopoietic aging.

While numerous NFκB system perturbations have shown its importance in hematopoiesis, they each have critical limitations.^15^ Mice with IKK and NIK mutations show bone marrow failure, but importantly, both IKK and NIK have pleiotropic targets outside the NFκB pathway.^22,29,30^ *Nfkb1*^-/-^ mice display features of premature aging, however, the biological consequences may be due to dysregulated NFκB, type I interferon, or dysregulated MAP kinase signaling.^15,31^ *Relb*^-/-^ mice have a complex inflammatory, autoimmune and immunodeficiency phenotype that is so severe that they are not suitable for aging studies.^32,33^ In contrast, NFκB activation achieved by reducing IκB-mediated sequestration of NFκB dimers in the cytoplasm is a direct and NFκB system-specific approach. Knock out mice for *Nfkbia* encoding IκBα show granulocytosis, but die shortly after birth, while *Nfkbia*^+/-^ heterozygous mice show no phenotype due to compensation by other IκB family members,^34^ limiting the utility of these mice for *in vivo* studies.

Here, we present a novel genetic model of inflammatory NFκB dysregulation termed IκB^−^ (*Nfkbia^+/-^Nfkbib^-/-^Nfkbie^-/-^*) in order to study the causal relationship between inflammation, myeloid bias, HSC accumulation, and HSC functional impairment seen in hematopoietic aging. We demonstrate the utility of the model as young IκB^−^ mice display myeloid bias and a rich inflammatory bone marrow milieu, similar to aged WT mice. Transplantation experiments in conjunction with epigenomic and transcriptomic analyses revealed inflammatory milieu-directed myeloid bias that is separable from HSC-intrinsic NFκB activity, quiescence and functional impairment. Together, the findings presented here delineate separate roles for inflammatory NFκB signaling in the bone marrow milieu and within HSCs themselves in driving different aspects of aging-associated hematopoietic dysregulation.

## RESULTS

### Experimentally elevated RelA drives myeloid-biased hematopoiesis similar to aging

In order to assess NFκB control in hematopoietic progenitors of aged mice, we used an mVenus-RelA reporter mouse previously generated in our laboratory.^35^ In this mouse, mVenus fluorescent signal indicates the abundance of RelA protein, which is the NFκB subunit most critical for inflammatory responses.^19,36^ In young mice (2-3 months) we found that hematopoietic stem and progenitor cell (HSPC) subsets, as phenotypically delineated by Pietras and colleagues^8^ (**Figure S1A**), vary in NFκB RelA protein abundance, with megakaryocyte-erythroid biased myeloid biased multipotent progenitor 2 (MPP2) cells showing the highest expression, and the lowest seen in short-term hematopoietic stem cells (ST-HSCs, also proposed to represent a multipotent progenitor rather than self-renewing HSC population^37^) (**Figures 1A-B and S1B**). Upon aging (18-22 months), RelA protein was increased in long-term hematopoietic stem cell (LT-HSC), ST-HSC, myeloid-biased MPP3 and lymphoid-biased MPP4 subsets (**Figures 1A, S1B-C**), consistent with a prior report.^18^

**Figure 1.**
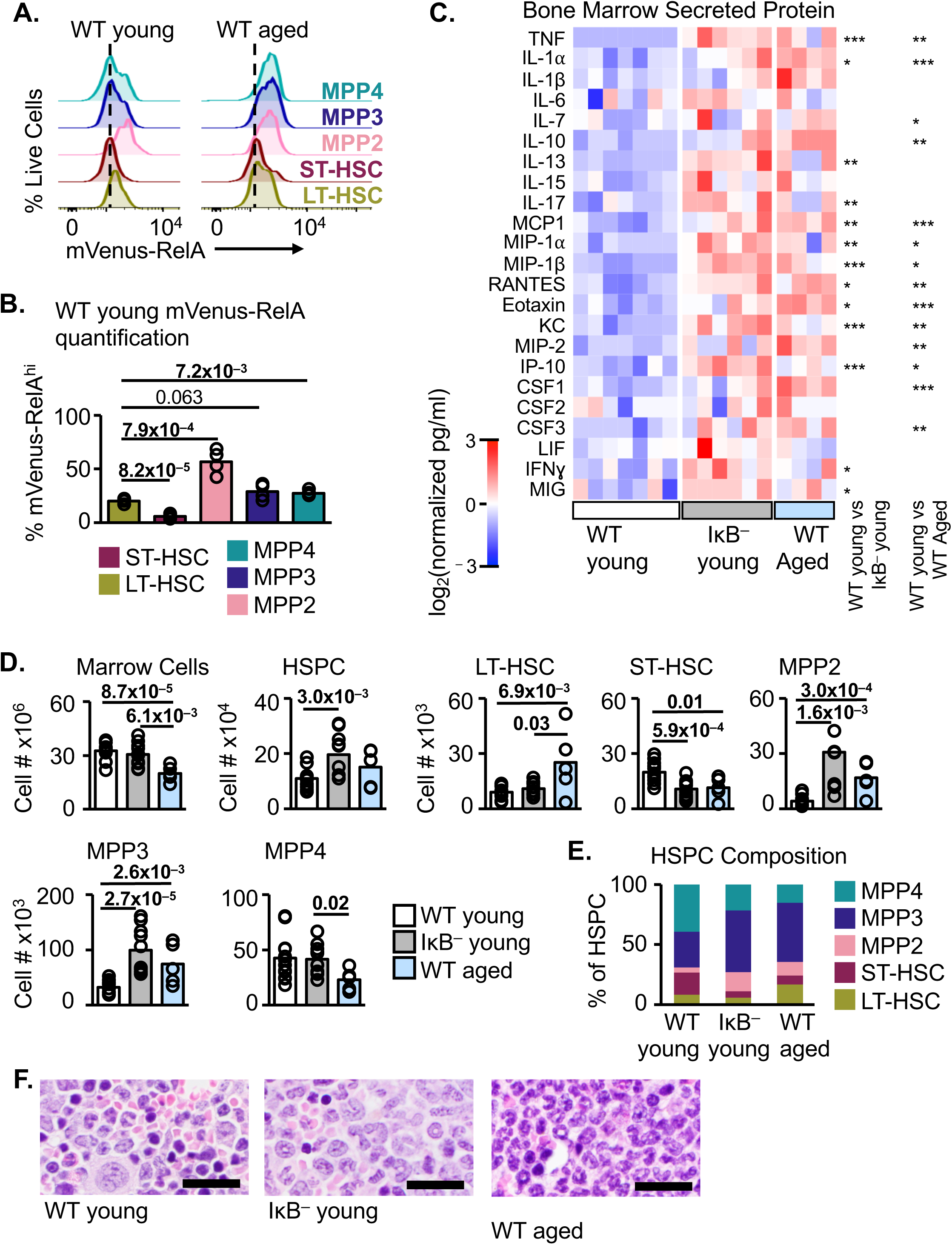
IκB^−^ mouse model of NFκB dysregulation has an inflamed bone marrow and myeloid biased hematopoiesis, similar to aged mice. **A.** mVenus-RelA fluorescence normalized to cell counts in the indicated HSPC subsets; representative of n=4-5. **B.** Quantification of RelA^hi^ young WT cells; statistics compare LT-HSC to other subsets. **C.** Secreted protein in bone marrow supernatant measured by Luminex, normalized to mean of WT young samples, per assay plate; n=4-7. **D.** Total bone marrow, total HSPC, and HSPC subset cell numbers by flow cytometry; n=5-12. **E.** Bone marrow HSPC subsets (as in D), displayed as a percent of total HSPCs; n=5-12. **F.** Sternum bone marrow histology, H&E; scale bars are 20μm; representative of n=3-5. Statistics calculated with unpaired students two-tailed t-test *p<0.05, **p<0.01, ***p<0.001. See also Figure S1.

To test whether elevated NFκB activity may be a sufficient driver of the aging-associated phenotypes of hematopoietic myeloid bias, HSC functional impairment, and HSC accumulation, we bred mice harboring knock-out alleles for genes encoding Inhibitor of κB (IκB) family members resulting in a compound IκB-deficient strain termed IκB^−^ (*Nfkbia*^+/-^*Nfkbib*^-/-^*Nfkbie*^-/-^).^34,38,39^ The combination of these genetic perturbations result in reduced cytoplasmic sequestration of NFκB dimers, leading to increased nuclear presence of NFκB and target gene expression. Accordingly, young IκB^−^ mice showed significant increases in soluble bone marrow inflammatory mediators over young WT controls. Indeed, the cytokine profile was found to be remarkably similar to the bone marrow inflammatory milieu observed in aged WT mice (**Figure 1C**).

Total bone marrow cellularity was similar between young IκB^−^ mice and young WT controls, however, the total number of HSPCs was increased in IκB^−^ (**Figure 1D**). This was predominantly due to increased abundance of megakaryocyte/erythroid-biased MPP2s and myeloid-biased MPP3s, without change in the number of lymphoid-biased MPP4s (**Figure 1D**). ST-HSCs were significantly reduced in IκB^−^ mice while LT-HSC abundance was unchanged when compared to young WT controls (**Figure 1D**). These alterations in HSPC subsets were appreciated in both the absolute numbers of cells in the bone marrow (**Figure 1D**) and when compared as a percent composition of the entire HSPC compartment (**Figure 1E**). Aged mice showed a similar increase in MPP2 and MPP3 subsets as IκB^−^, and additional changes were noted including a reduction in total bone marrow cellularity, decrease in MPP4s, and increase in LT-HSCs (**Figures 1D-E, S1D**), consistent with established literature in aged mice.^1,2^ Within the LT-HSC compartment, cell surface expression of CD150 has been associated with lineage biases.^40,41^ CD150^high^ myeloid-biased HSCs (myHSCs) accumulate with aging,^40,41^ which we observed our aged murine cohort (**Figure S1D**). However, IκB^−^ mice showed variable proportions of myHSCs vs. CD150^low^ lymphoid-biased (lyHSCs), without overall increase in myHSCs detected (**Figure S1D**), consistent with the overall lack of HSC accumulation noted in this model.

Histologic evaluation of the bone marrow revealed mature granulocytosis in both young IκB^−^ and aged WT mice compared with young WT controls (**Figure 1F**). In the peripheral blood, young IκB^−^ mice showed a reduction in circulating B-cells (further explored in Lin et al., in preparation) and an increase in myeloid cells (granulocytes and myelomonocytic cells); no difference in T-cell abundances was detected (**Figure S1E-F**). These findings were similar, though more pronounced, than those observed in aged WT mice (**Figure S1F**), and indicate myeloid bias among mature immune effector cells in the peripheral circulation. Together, these results in peripheral blood and bone marrow indicate that experimentally driving ubiquitous proinflammatory NFκB signaling is sufficient for myeloid-biased hematopoiesis as seen in aged mice and humans, but does not lead to aging-associated HSC accumulation.

### Hematopoietic-cell-extrinsic NF**κ**B activity is sufficient to drive myeloid bias

The striking abundance of soluble inflammatory cytokines and chemokines in IκB^−^ and aged bone marrow (**Figure 1C**) raises the possibility that they may be responsible for driving myeloid bias. These soluble factors may be produced by either hematopoietic cells including HSPCs themselves, or by non-hemopoietic cells. To distinguish between these possibilities, we generated bone marrow chimeric mice by transplanting WT donor bone marrow into IκB^−^ recipients or WT controls (**Figure 2A**). This strategy allowed us to measure how elevated NFκB activity in the radio-resistant, hematopoietic-cell-extrinsic compartment affects WT donor cells. Transplant recipients of both genotypes showed excellent chimerism 15 weeks post-transplant (>97%; **Figures 2B and S2A**), a time point when donor HSCs are expected to show stable engraftment, and complete hematopoietic reconstitution.^42^ Experimental hematopoietic-extrinsic NFκB dysregulation in IκB^−^ recipients resulted in unchanged total bone marrow cellularity and total HSPC numbers between recipient genotypes (**Figure 2C**); however, myeloid-biased MPP3s were increased in WT donor cells from IκB^−^ recipients measured by absolute number (**Figure 2C**) and as a proportion of the entire HSPC compartment (**Figure 2D**), compared to WT recipient controls. Mature granulocytosis in the bone marrow was seen by histology (**Figure S2B**), and peripheral blood immunophenotyping showed increased myeloid cells (**Figure S2C**), indicating myeloid bias among mature immune effector cells. Thus, myeloid bias of stably-engrafted WT donor cells in the periphery and bone marrow of IκB^−^ recipients was similar to that observed in non-transplanted IκB^−^ mice (**Figures 1D-E and S1F**). In contrast, the increase in megakaryocyte/erythroid-biased MPP2s and decrease in ST-HSCs present in non-transplanted IκB^−^ mice (**Figure 1D-E**) were not seen after primary transplantation into IκB^−^ recipients (**Figure 2C-D**). Therefore, NFκB-activity restricted to the radio-resistant compartment is sufficient for myeloid bias.

**Figure 2.**
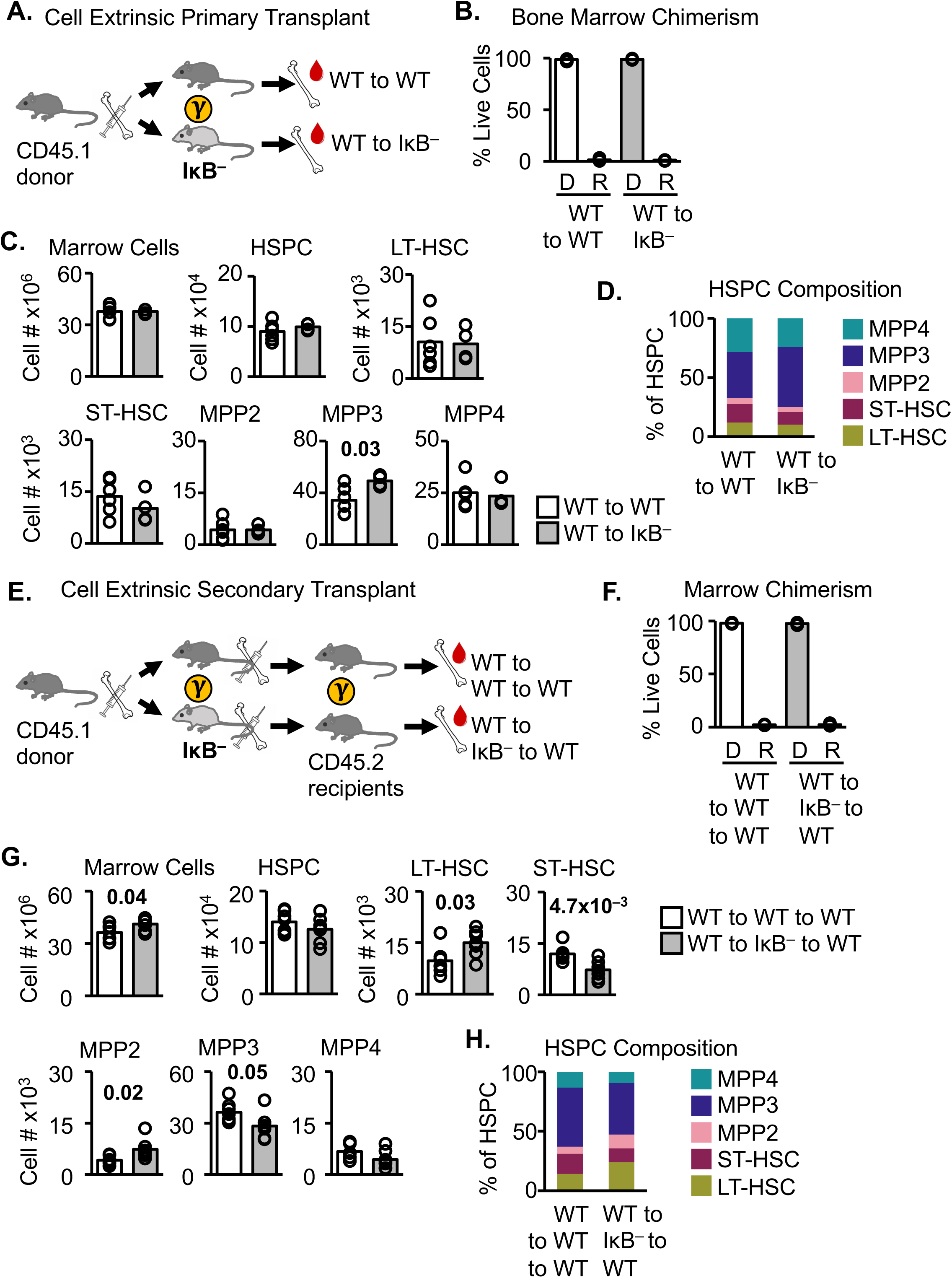
Hematopoietic cell-extrinsic inflammatory signals are sufficient to drive myeloid biased hematopoiesis. **A.** Primary cell-extrinsic transplant schematic. **B.** Bone marrow chimerism 15 weeks following primary transplant; Donor (D), Recipient (R). **C.** Donor CD45.1 bone marrow, HSPC, and HSPC subset cell numbers. **D.** Donor CD45.1 bone marrow HSPC subsets (as in C), displayed as a percent of total donor HSPCs. **E.** Secondary cell-extrinsic transplant schematic. **F.** Bone marrow chimerism 16 weeks following secondary transplant; Donor (D), Recipient (R). **G.** Donor CD45.1 bone marrow, HSPC, and HSPC subset cell numbers. **H.** Donor CD45.1 bone marrow HSPC subsets (as in G), displayed as a percent of total donor HSPCs. A-D n=4-6; E-H n=7. Statistics calculated with unpaired students two-tailed t-test. See also Figure S2.

We next tested the durability of milieu-directed myeloid bias upon return to a WT bone marrow milieu using secondary transplantation. Secondary transplantation of WT marrow conditioned in IκB^−^ primary recipients for 15 weeks into young WT secondary recipients (**Figure 2E**) led to a remarkable resolution of MPP3 bias after 16 weeks (**Figure 2F-H**). Together, these experiments demonstrate that HSC-extrinsic NFκB activity is sufficient to drive myeloid-biased hematopoiesis, but that NFκB-driven inflammatory signals in the milieu must be persistent to maintain progenitor myeloid bias. Furthermore, NFκB activity in the bone marrow milieu does not drive expansion of megakaryocyte/erythroid-biased progenitors, nor a reduction in HSCs, suggesting specificity for milieu-derived signals in promoting myeloid bias. Finally, these data demonstrate that HSPC myeloid bias is reversible upon engraftment of HSCs into a non-dysregulated bone marrow milieu.

### The inflamed bone marrow milieu directs epigenomic reprogramming of HSCs

We next asked whether epigenomic changes may underly the HSPC myeloid bias induced in WT cells by the inflamed bone marrow milieu of IκB^−^ mice. We performed Assay for Transposase Accessible Chromatin with sequencing (ATAC-seq)^43^ in flow cytometry-sorted LT-HSC (HSC), MPP2, MPP3 and MPP4 populations from WT-to-WT or WT-to-IκB^−^ bone marrow chimeric mice (**Figure S3A**), identifying 13910 consensus peaks of chromatin accessibility among 3 biological replicates. We confirmed that such peaks were appropriately larger for regions expected to be open in all samples (e.g. *Gapdh*) than for regions expected to be closed in all samples (e.g. *Pbpb*), and showed appropriate cell-type specificity (e.g. *Pf4*, larger in megakaryocyte/erythroid-primed MPP2s) (**Figure 3A**).

**Figure 3.**
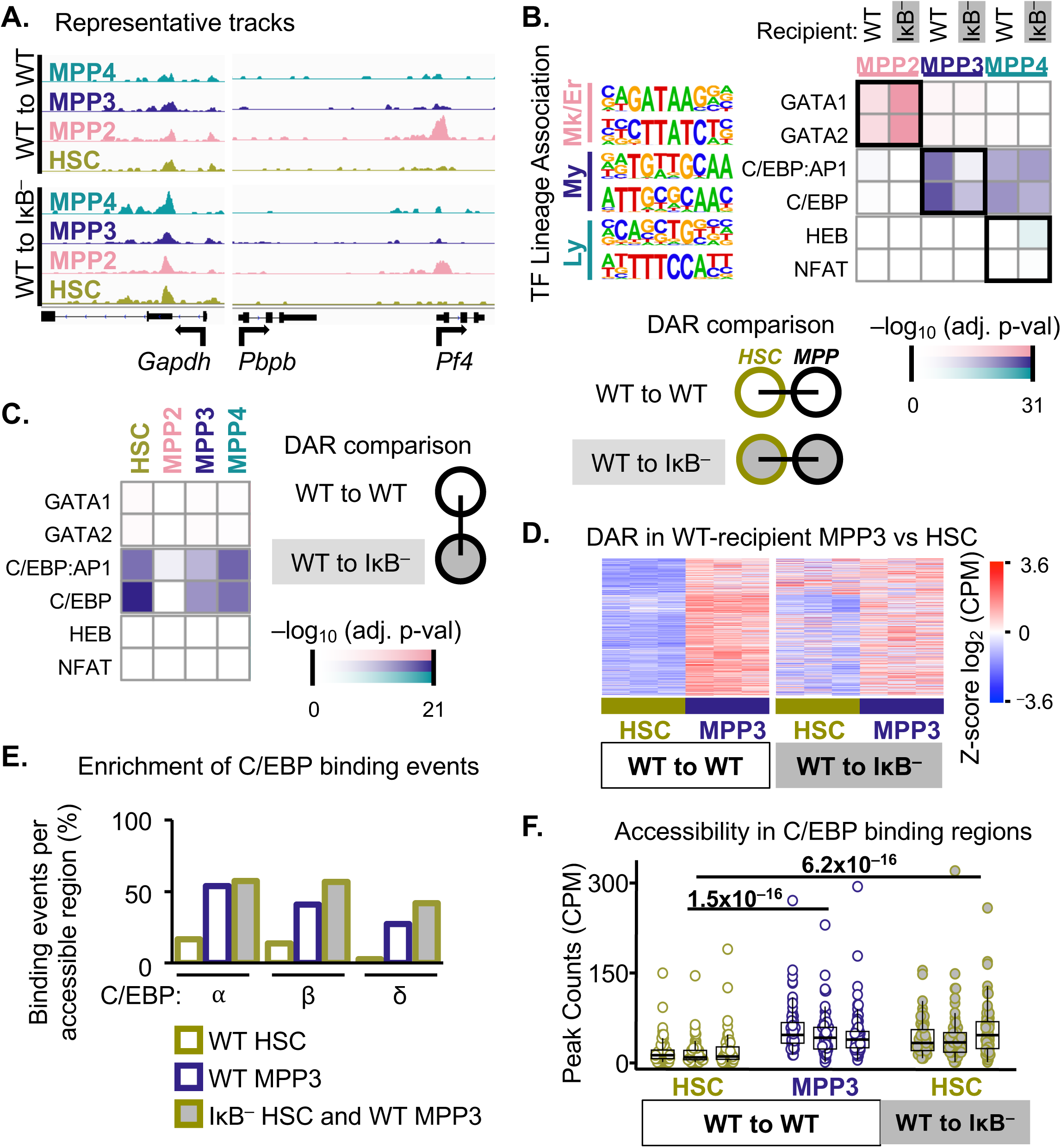
In an inflammatory milieu, HSC undergo chromatin reorganization that results in increased accessibility for myeloid-priming C/EBP transcription factors. **A.** Representative tracks of peaks that correspond to chromatin accessibility for regions expected to be open in all samples (*Gapdh*), closed in all samples (*Pbpb*) or population specific (*Pf4*); n=3 biological replicates. **B.** Transcription factor (TF) motif enrichment among differentially accessible chromatin regions (DAR) that are more accessible in the specified MPP population vs. HSC, separately for each recipient. **C.** TF motif enrichment in DAR more accessible in IκB^−^ recipient vs. WT recipient, for each population. **D.** Z-score of CPM normalized peak counts in the 595 DAR that are more accessible in WT-recipient MPP3 than in WT-recipient HSC, shown for all 3 biological replicates. **E.** Overlap between published chromatin immunoprecipitation C/EBP binding events and ATAC-seq DAR for regions accessible in WT-recipient HSC but not WT-recipient MPP3 (WT HSC), in WT-recipient MPP3 but not WT-recipient HSC (WT MPP3), or in WT-recipient MPP3 and IκB^−^-recipient HSC, but not WT-recipient HSC (IκB^−^ HSC and WT MPP3). **F.** CPM normalized peak counts in “IκB^−^ HSC and WT MPP3” regions that overlap with any C/EBP binding event, for the indicated populations and all replicates. Statistics in B and C reported as BH adjusted p-values calculated via Homer ZOOPS scoring; statistics in F calculated with Wilcoxon Rank Sum test. See also Figure S3.

To identify differentiation-associated changes in chromatin accessibility for each progenitor lineage, we used differential accessibility analysis between each MPP subset vs. HSCs, performed separately for cells from WT and IκB^−^ recipients (**Figure S3B**). In these differentially accessibly regions (DARs), we then assessed TF motif enrichment in megakaryocyte/erythroid-biased (Mk/Er) MPP2s, myeloid-biased (My) MPP3s, and lymphoid-biased (Ly) MPP4s (**Figure 3B**).^44–48^ In contrast to WT-recipient controls, IκB^−^ recipient myeloid-associated MPP3 vs. HSC DARs showed little enrichment for CCAAT/enhancer-binding protein (C/EBP) motifs (**Figure 3B**), which was surprising given the myeloid-biased phenotype in IκB^−^ recipient mice (Figure 2) and C/EBP’s prominent role in myeloid differentiation.^45^ While this analysis related DARs of MPPs vs HSCs within the same recipient, we next directly compared IκB^−^ vs. WT recipient DARs for each HSPC population to identify changes in chromatin accessibility driven by the inflammatory milieu (**Figure S3C, Figure 3C**). This analysis revealed a dramatic enrichment of C/EBP motifs among IκB^−^-recipient HSCs, and moderate enrichments in MPP3 and MPP4 populations (**Figure 3C**). A similar pattern was noted for PU.1 (**Figure S3D**), which instructs both myeloid and early lymphoid programs,^49^ but not for megakaryocyte/erythroid-associated nor lymphoid-specific TF motifs (**Figure 3C**).

We next identified specific chromatin regions associated with the myeloid lineage that become prematurely accessible in WT HSCs transplanted into an inflamed IκB^−^ milieu. Among 595 myeloid-specific DARs in WT recipients (defined as those more accessible in WT-recipient MPP3s vs. HSCs), a distinct cluster of 115 DARs showed higher peak counts in IκB^−^-recipient HSCs than controls (**Figure 3D**). C/EBPα and C/EBPβ have well-established roles in steady-state and emergency myelopoiesis, respectively, and C/EBPδ is implicated in granulocyte specification and function.^45,46^ Furthermore, C/EBPα and C/EBPβ are pioneer factors that can open heterochromatin, thereby facilitating a change in transcriptional programs, either independently or synergistically with PU.1.^50^ Thus, we asked whether these myeloid-associated DARs may overlap with *bona fide* C/EBP binding events, which were defined from publicly available chromatin immunoprecipitation with sequencing (ChIP-seq) datasets of differentiated myeloid cells.^51–53^ Indeed, we found that the 115 myeloid regions prematurely accessible in IκB^−^-recipient HSCs showed a high overlap with C/EBP binding events, as did total WT-recipient myeloid-specific MPP3 vs. HSC DARs (595), while negative control WT-recipient HSC-specific DARs (434) did not (**Figure 3E**), confirming that newly accessible regions in HSCs from an IκB^−^ milieu are enriched in *bona fide* C/EBP binding sites. Regions with high C/EBP overlap also showed higher peak counts, indicated increased chromatin accessibility at these C/EPB binding sites (**Figure 3F**). Together, these findings identify inappropriate or premature accessibility for C/EBPs and their binding in HSCs conditioned in an inflammatory IκB^−^ milieu as a form of epigenomic reprogramming.

### The inflamed bone marrow milieu redirects the gene expression program of HSCs

Next, we asked whether WT HSPCs from an inflammatory IκB^−^ milieu may also show transcriptional features of myeloid bias. We performed single-cell RNA sequencing (scRNA-seq) on flow cytometry-sorted HSPC from WT-to-WT or WT-to-IκB^−^ bone marrow chimeric mice (**Figure S4A**). To identify HSPCs that relate to the individually sorted subsets in our ATAC-seq analysis, we developed a logistic-regression-based cell labeling model. This model generated HSPC subset labels from bulk-sorted LT-HSC, ST-HSC, MPP2, MPP3 and MPP4 microarray data, which were then filtered by an independent bulk-sorted scRNA-seq dataset, both of which used the same phenotypic cell surface markers as our ATAC-seq sorting strategy.^8,54^ In test scRNA-data, cell-type labels were assigned as one of the five HSPC cell types that showed the highest transcriptomic correspondence between test and reference data (**Figure S5**). Labeling our cell-extrinsic transplant scRNA-seq data resulted in more MPP3-labeled cells from IκB^−^-recipients than WT-recipient controls (**Figures 4A-B, Figures S4B-D**), consistent with flow cytometry phenotyping. We first evaluated the expression of genes nearest the accessible C/EBP binding events identified in Figures 3D-F. These transcripts were increased among the differentially expressed genes (DEG) from positive control WT-recipient MPP3s vs. WT-recipient LT-HSCs as expected (**Figure 4C**, x-axis), and were also higher in LT-HSCs from IκB^−^-recipients vs. control LT-HSCs (**Figure 4C**, y-axis), confirming increased transcription of most C/EBP target genes in HSCs conditioned in an inflamed environment. We next sought to perform similar transcript abundance analysis while retaining single cell resolution. To do so, we calculated rank-based single cell gene set enrichment scores,^55^ yielding a scoring metric that indicates how highly expressed a set of genes is within each cell. For ATAC-accessible C/EBP target genes, single cell gene set scores showed an increase in C/EBP target gene score, consistent with the DEG-based analysis (**Figure 4D**). These findings confirmed increased transcript abundance for genes in C/EBP binding regions that were prematurely accessible in HSCs conditioned in an inflamed IκB^−^ milieu.

**Figure 4.**
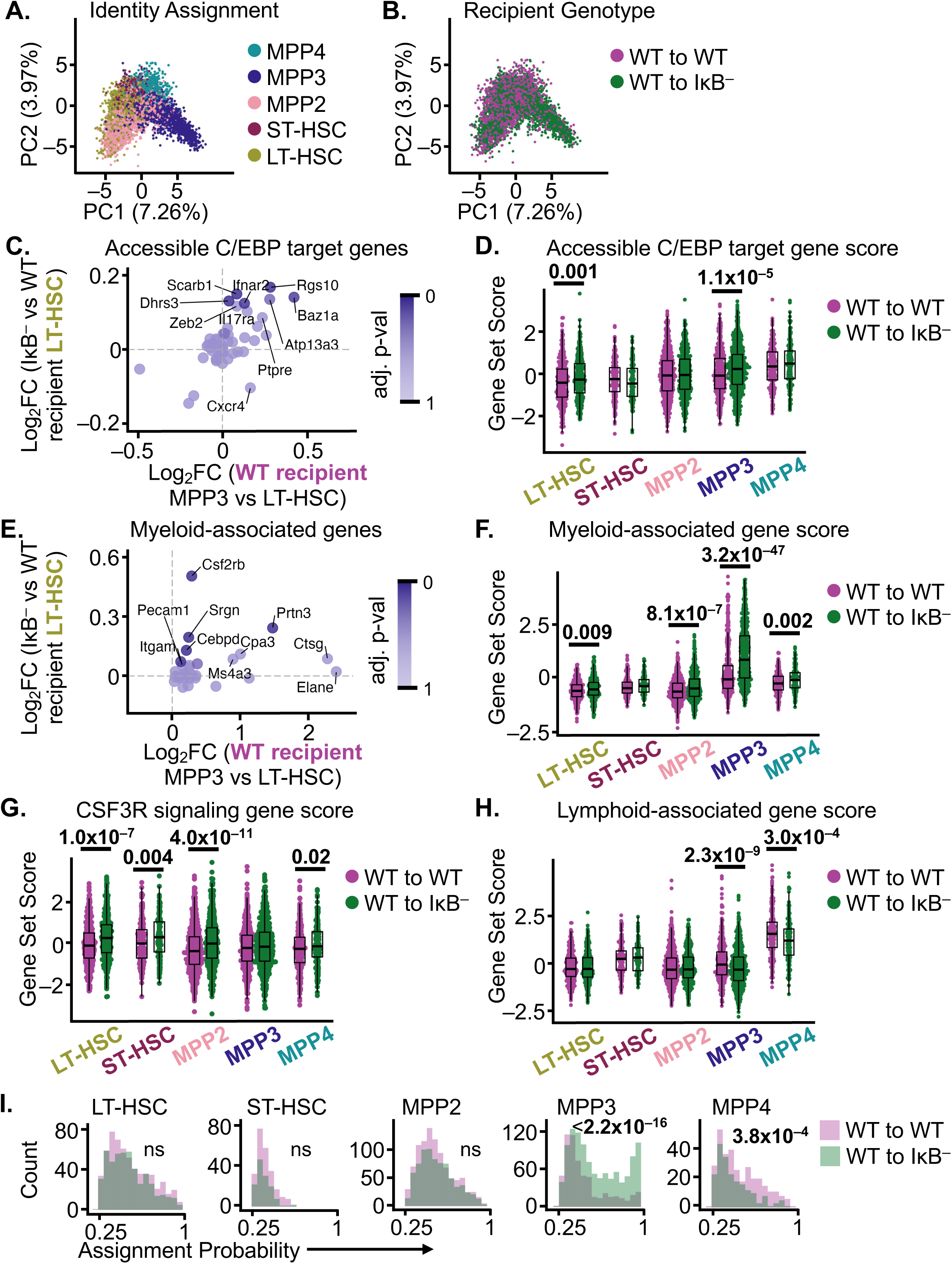
HSPC in an inflammatory milieu show a myeloid-biased transcriptional state. scRNA-sequenced cells from cell-extrinsic transplants (as in Figure 2A-D) labeled by cell identity assignment (**A**) or by recipient genotype (**B**), plotted in PCA space defined by expression of 115 genes with known functions in hematopoiesis. Expression of genes corresponding to “IκB^−^ HSC and WT MPP3” DARs that overlap with C/EBP binding events (as in Fig 3E-F), expressed as population-level differentially expressed genes (DEG); x-axis shows positive control WT-recipient MPP3s vs. WT-recipient LT-HSCs; y-axis experimental LT-HSCs from IκB^−^-recipients vs. control LT-HSCs (**C**) or expressed as rank-based single cell gene set enrichment score (**D**). Expression analysis of 31 myeloid-associated genes^56^ by population DEG; x-axis shows positive control WT-recipient MPP3s vs. WT-recipient LT-HSCs; y-axis experimental LT-HSCs from IκB^−^-recipients vs. control LT-HSCs (**E**) and single cell gene set score (**F**). Single cell set scores for CSF3R response genes^57^ (**G**) and lymphoid-associated genes^58^ (**H**). **I.** Cell assignment probabilities among each cell subset identity assignment. Statistics calculated with Wilcoxon Rank Sum test (A-G) or KS test of the distribution (I). BH-adjusted p-values in C and E correspond to y-axis DEG. See also Figures S4-S5 and Table S1.

We next evaluated broader transcriptional features of myeloid bias. Expression of 31 myeloid lineage- and maturation-associated genes encoding myeloid TFs, growth factor receptors, and functional proteins^56^ (**Table S1**) were increased in LT-HSC IκB^−^ recipients compared to WT recipients in both DEG (**Figure 4E**) and single-cell gene set scoring approaches (**Figure 4F**). Furthermore, myeloid-associated genes showed higher single cell scores among IκB^−^ recipient MPPs than WT recipient controls (**Figure 4F**). To confirm our findings with an orthogonal, non-overlapping gene set, we identified a set of 49 CSF3R response genes from the NCI Cytosig database,^57^ including *Ctsb*, *Nampt*, and *Timp1* (**Table S1**). Similar to myeloid-associated genes, single-cell gene set scores were higher for CSF3R-response genes in HSCs and non-MPP3 progenitors from IκB^−^ recipients (**Figure 4G**). Lymphoid-associated transcripts were also evaluated using a gene set of 11 lymphoid lineage-associated genes^58^ including *Dntt*, *Rag1*, and *Ikzf1* (**Table S1**). Lymphoid gene set scores were unchanged in HSCs, and were reduced in MPP3s and MPP4s from IκB^−^ recipients compared to WT recipients (**Figure 4H**). While together these three myeloid-associated, CSF3R-responsive and lymphoid-associated gene sets contain <100 genes, our cell-type labeling model incorporates expression data of 7563 genes to classify HSPCs and resolve lineage-associated transcriptional features among transcriptomically similar stem and progenitor cells. Thus, we next examined the distributions of cell type assignment probabilities from our labeling model. This revealed that among MPP3-labeled cells, IκB^−^-recipient cells had higher MPP3 assignment probabilities than WT recipient controls (**Figure 4I**). Conversely, among MPP4-labeled cells, IκB^−^-recipient MPP4s instead had lower MPP4 assignment probabilities (**Figure 4I**). Together, these findings identify transcriptional myeloid bias in LT-HSCs and MPPs conditioned in the inflammatory IκB^−^ milieu.

### Milieu-directed transcriptional bias of HSCs in aged mice and humans

To determine whether transcriptional signatures identified in HSCs from an inflammatory milieu are present in samples from naturally aged individuals, we evaluated publicly available scRNA-seq datasets from murine HSPCs and human CD34-positive cells.^59–62^ We applied our cell labeling model to each dataset to ensure fair comparisons between them (**Figure 5A-B, and Figure S6A**), and confirmed suitability of our labeling method for human data using species-specific reference genes^63^ (**Figure S6B**). In the absence of chromatin accessibility data, we computationally inferred TF activity from scRNA-seq data using the decoupleR tool that leverages independent gene regulatory and transcriptomic databases.^64^ Comparing the scRNA-seq DEGs of aged versus young LT-HSCs from mice and humans, we found a combination of increased myeloid (My) TF activity and reduced lymphoid (Ly) TF activity,^65,66^ pointing to an overall myeloid-biased TF repertoire (**Figure 5C**). To assess myeloid potentiation and lymphoid signature decline in a single metric, a bias score was calculated as the difference between the CSF3R-response or myeloid-associated single-cell gene set enrichment scores, and the lymphoid-associated single cell gene set enrichment score. The resultant CSF3R-bias scores were strongly increased in LT-HSC from all aged datasets (**Figure 5D**), and myeloid-bias scores were increased in 3 of 4 datasets (**Figure 5E**). Slight differences between these scores may reflect early vs. later events in myeloid commitment, as LT-HSCs from cell-extrinsic transplantation experiments showed higher scores for CSF3R responses than for myeloid-associated genes (**Figure 4F-G**). These findings indicate transcriptional myeloid bias in aged HSCs from mice and humans, similar to that identified in murine HSCs from an inflamed IκB^−^ milieu.

**Figure 5.**
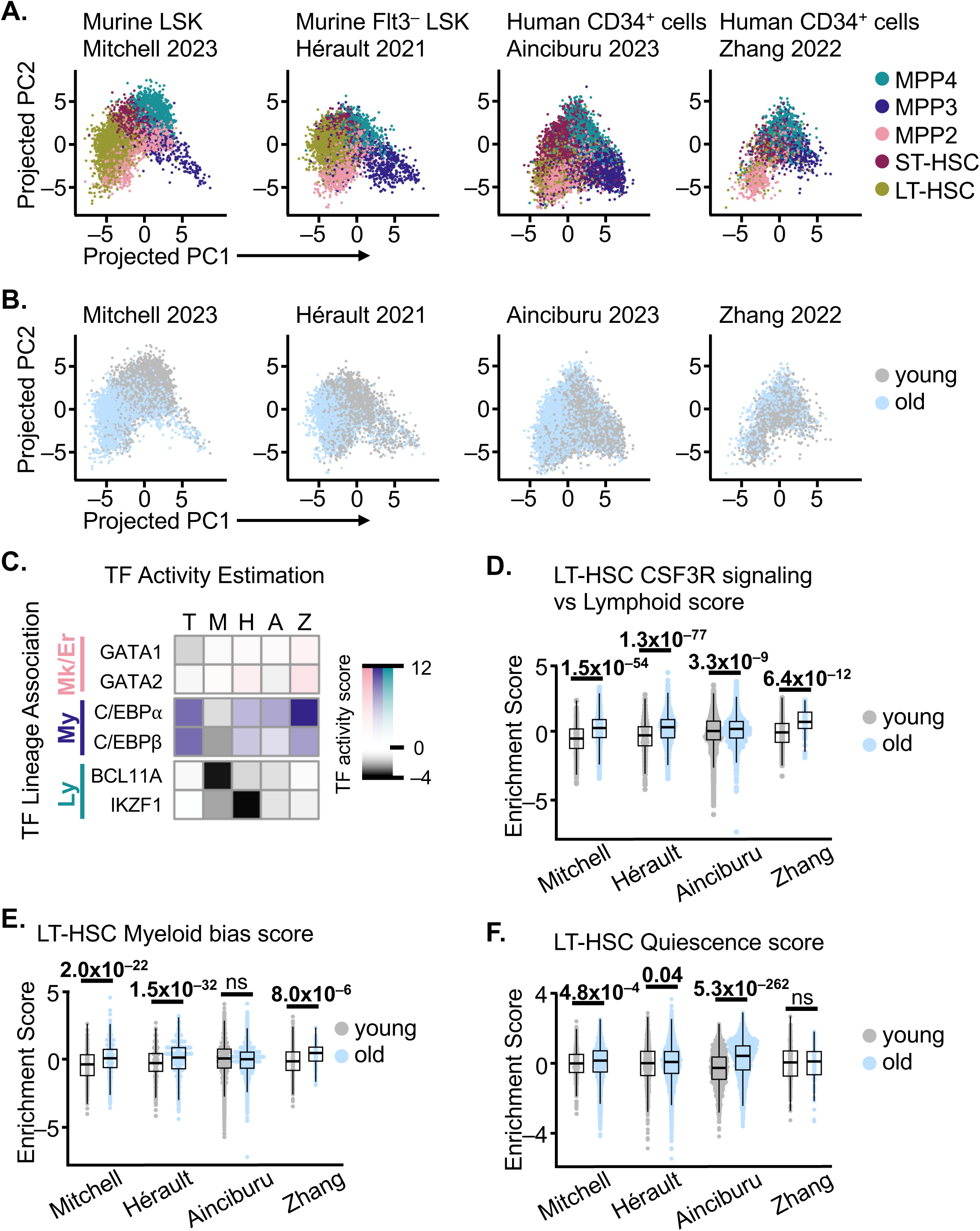
Aged HSC from mice and humans show transcriptional myeloid bias and quiescence signatures. scRNA-sequenced cells from previously published datasets projected onto PCA space from the transplant dataset (Figure 4A-B), labeled by cell identity assignment (**A**) or age (**B**). **C.** TF activity estimation^64^ in transplant (T), Mitchell 2023 (M), Hérault 2021 (H), Ainciburu 2023 (A) and Zhang 2023 (Z) datasets. CSF3R bias (CSF3R response -lymphoid-associated genes) (**D**), myeloid bias (myeloid-associated -lymphoid-associated) genes (**E**), and quiescence gene^68^ set score (**F**) among LT-HSC. Statistics calculated with Wilcoxon Rank Sum test. See also Figure S6 and Table S1.

### Aged HSCs show transcriptomic signatures of quiescence or myeloid bias, but not both

HSCs remain in quiescence, a reversible G0 state of dormancy, until receiving an activation cue to enter the cell cycle.^4^ This results in a long-lived pool of HSCs that can maintain multilineage hematopoietic potential throughout the life span, while progenitor proliferation contributes to the bulk of hematopoietic output.^1^ Importantly, quiescence is now recognized as a spectrum, rather than a binary on/off state, and increased quiescence has been correlated with the HSC accumulation observed in aging, though the causal connection remains unclear.^1,67^ In the scRNAseq datasets we evaluated, aged HSCs were more abundant than young in all 3 datasets that had no further cell sorting or selection (**Figure S6C**, Mitchell 2023, Hérault 2021 and Ainciburu 2023), consistent with the literature^5,6^ and our own data (**Figure 1E**). To explore the depth of quiescence in aged HSCs, we used 73 genes with well-established roles in quiescence, including *Ccnd3*, *Ezh1*, and *Cdc42* (**Table S1**), to calculate single cell gene set scores.^68–70^ Indeed, aged LT-HSCs showed higher single cell quiescence gene set scores than young for all datasets with HSC accumulation (**Figure 5F**). Thus, both transcriptional myeloid bias and increased quiescence signatures are consistently observed in aged HSCs from mice and humans.

We next leveraged the single-cell resolution of these data to examine the relationship between transcriptomic myeloid bias and quiescence states. On a per-cell basis, we surprisingly observed no correlation between myeloid bias and quiescence scores (**Figure 6A**). We then asked how highly quiescent HSCs differ from myeloid-biased HSCs. To do so, we performed a DEG analysis between aged LT-HSCs that have high quiescence scores (top 25^th^ percentile, pooled among datasets to increase statistical power) vs. those with high myeloid bias scores (**Figure 6B**). In aged LT-HSCs with high quiescence scores, there was significant downregulation in terms for cell cycle and mitotic division, as expected (**Table S2**). The top upregulated gene ontology (GO) terms were for cell signaling, migration and morphogenesis, consistent with known cytoskeletal changes that occur in aging^71^ (**Table S3**). Intriguingly, the GO Immune Response term was also significantly enriched (BH-adjusted p-value 1.9×10^−9^). Thus, the aged HSC pool shows heterogeneity with distinct transcriptional programs of myeloid bias and quiescence.

**Figure 6.**
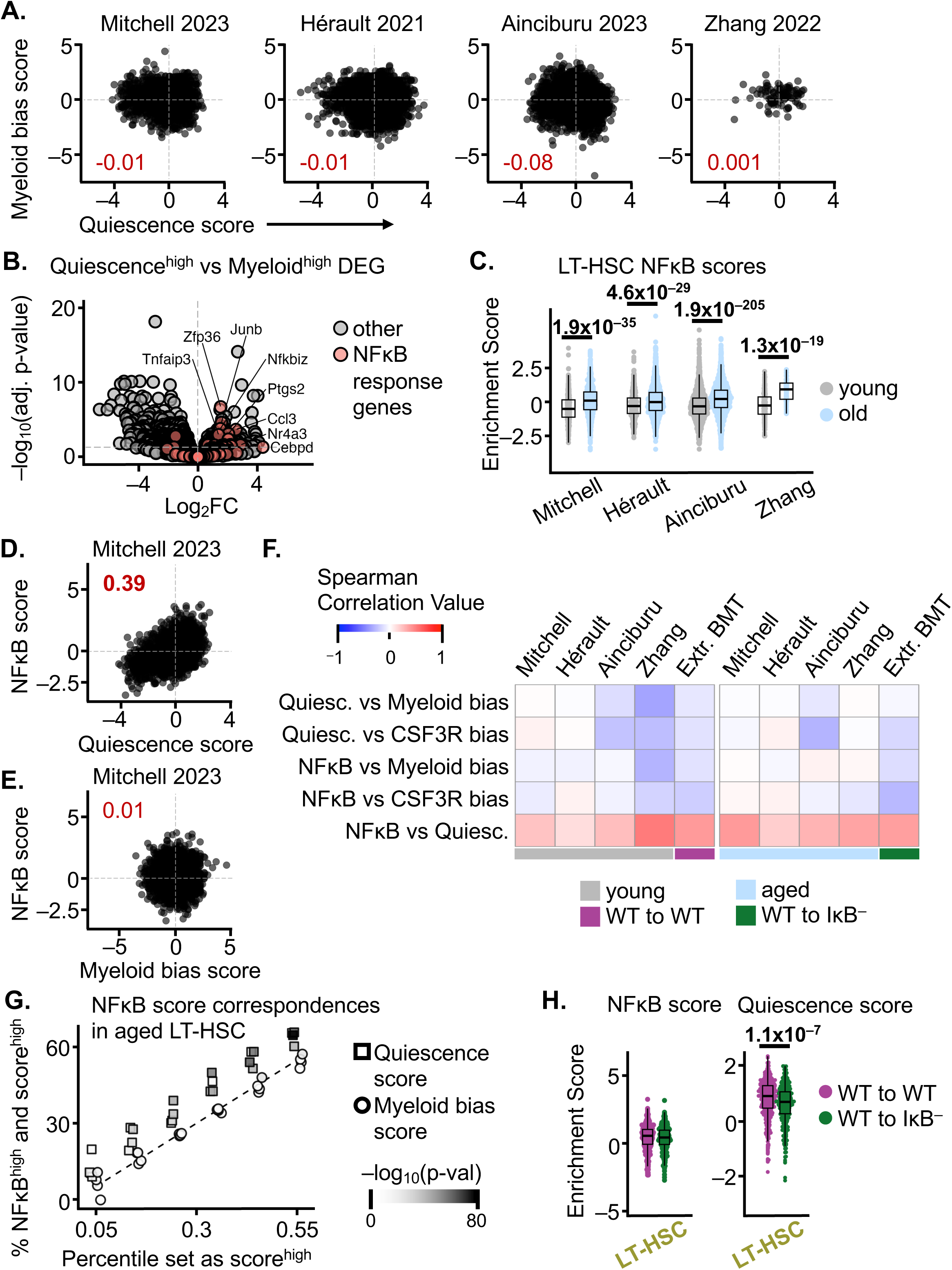
In HSCs, NFκB signatures correlate with quiescence but not myeloid bias. **A.** LT-HSC single cell correspondence between quiescence and myeloid versus lymphoid gene set scores; insets indicate Spearman correlation value. **B.** DEG between cells that are in the top 25^th^ percentile for quiescence score (“Quiescence^high^”) versus top 25^th^ percentile for myeloid bias (“Myeloid^high^”, myeloid versus lymphoid score), combined for all old LT-HSC across datasets. BH-adjusted p-values. **C.** NFκB response gene scores among LT-HSC in aged datasets. **D.** LT-HSC single cell correspondence between quiescence and NFκB response gene set scores; insets indicate Spearman correlation value. **E.** LT-HSC single cell correspondence between NFκB response gene and myeloid bias scores; insets indicate Spearman correlation value. **F.** Spearman correlations between single cell gene scores, for all datasets and conditions. **G.** For aged LT-HSCs at specific percentile thresholds (x-axis), percent of cells that reach both NFκB^high^ and Quiescence^high^ (square), or both NFκB^high^ and Myeloid^high^ (circle) thresholds; the dashed line indicates the expected values if no relationship between NFκB score and the comparator score is observed. **H.** NFκB-response gene and quiescence scores among LT-HSC from transplant dataset. Statistics in C, F and G calculated with Wilcoxon Rank Sum test. See also Tables S1-S3.

### HSC-intrinsic NF**κ**B activity correlates with quiescence but not myeloid bias

As NFκB controls immune responses, we next examined whether known NFκB-response genes are differentially expressed in highly quiescent vs. myeloid-biased HSCs. Indeed, NFκB-response genes^72^ were 3.1-fold more abundant among the upregulated DEGs from highly quiescence HSCs (**Figure 6B**). Further, single-cell NFκB gene set enrichment scores using sets of 270 murine and 248 human NFκB-responsive genes^72^ including *Nfkbia*, *Ccl5*, and *Il1b* were increased in aged LT-HSCs compared to young controls in all datasets examined (**Figure 6C**). Two orthogonal approaches demonstrated a relationship between NFκB and quiescence scores in aged HSCs, but found no relationship between NFκB and myeloid bias scores, either by calculating Spearman correlation between single-cell scores (**Figure 6D-F**), or by determining the percentage overlap between high-scoring cells (**Figure 6G**). Notably, the correlation between NFκB and quiescence was consistent regardless of HSC age (**Figure 6F**), suggesting a consistent biological relationship that is actually independent of aging, though both signatures are potentiated in HSCs from aged individuals.

We next asked whether myeloid-biased HSCs from an inflamed IκB^−^ milieu may show similar relationships between transcriptional myeloid bias, NFκB activity, and quiescence as seen in aged HSCs, utilizing scRNA-seq data from our cell-extrinsic IκB^−^ transplantation studies. Intriguingly, LT-HSCs from IκB^−^ recipients showed unchanged NFκB scores compared to controls (**Figure 6H**, left) despite their myeloid-biased transcriptomic and epigenomic states (**Figures 3** and **4**). This distinction underscores the separation of HSC-intrinsic NFκB activity and milieu-directed myeloid bias. Furthermore, LT-HSCs from IκB^−^-recipients (**Figure 6H**, right) showed reduced quiescence scores, consistent with the lack of HSC accumulation observed in these mice. Positive per-cell correlations between NFκB and quiescence, but not between NFκB and myeloid bias/CSF3R bias, were similar in cell-extrinsic transplantation experiments to those seen in young and aged HSCs (**Figure 6F**). These findings demonstrate that the transcriptional programs supporting NFκB activity, myeloid bias, and quiescence are all increased in HSCs upon aging. However, on a per-cell basis, HSC-intrinsic expression of NFκB target genes is associated with higher quiescence signatures, but not with myeloid bias. The per-cell relationships between these transcriptional programs are conserved in murine and human HSCs across the age-span.

### Cell-intrinsic NF**κ**B activity drives functional HSC impairment

In addition to the close transcriptional relationship between NFκB activity and quiescence, we also observed that NFκB RelA protein in mVenus-RelA reporter mice is more abundant in quiescent LT-HSCs than in more proliferative ST-HSCs (**Figure 1A-B**), further suggesting a functional relationship between NFκB and quiescence. To explore this possibility, we next tested whether experimentally elevating NFκB within HSCs impacts their functional reconstitution capacity using a transplantation approach. NFκB activity alters expression of *Cxcr4* (**Figure 4C**),^73^ a critical chemokine for HSC homing. Thus, competitive transplantation between IκB^−^ HSCs that harbor experimentally elevated NFκB vs. WT controls may show early differences in niche homing and occupancy that would overshadow quiescence-mediated phenotypes. Thus, we performed IκB^−^ cell-intrinsic transplantation experiments in a non-competitive manner to ensure that observations of long-term engraftment and bone marrow reconstitution after 4 months, when donor hematopoietic cells are derived from engrafted HSCs,^42^ are not attributable to differences in early competitive HSC homing.

Primary bone marrow transplantation performed with IκB^−^ donor cells into WT recipients revealed reduced engraftment compared to WT donor controls (peripheral blood chimerism 94% in WT donor vs. 69% in IκB^−^ donor; bone marrow chimerism 99% in WT donor vs. 88% in IκB^−^ donor) after 16 weeks (**Figures S7A and 7A-B**). Peripheral blood immunophenotyping confirmed that the marked reduction in circulating B-cells seen in IκB^−^ mice is mediated through a cell-intrinsic mechanism (**Figure S7B**). In the bone marrow, total cellularity did not differ between mice receiving WT vs. IκB^−^ donor marrow. However, an increase in total HSPCs was noted in IκB^−^ donor marrow, driven by increases in MPP2s, MPP3s and MPP4s, and opposed by a significant decrease in ST-HSCs (in absolute numbers **Figure 7C**, and by percent of HSPCs, **Figure 7D**). In addition, LT-HSC numbers were significantly reduced in IκB^−^ donor marrow compared to WT controls (**Figure 7D**), a contrast with non-transplanted IκB^−^ mice (**Figure 1D**). Of note, HSPCs and maturing hematopoietic cells not only respond to, but also release proinflammatory cytokines and other paracrine signals.^74–79^ Thus, transplanted IκB^−^ hematopoietic cells may secrete mediators that promote an inflammatory milieu and/or remodel their niches. Therefore, the findings of HSPC expansion and ST-HSC reduction common to both IκB^−^ cell-intrinsic transplants and non-transplanted IκB^−^ mice may be attributable to either cell-intrinsic effects, or to responses to hematopoietic-derived inflammatory mediators. In contrast, the reduction in LT-HSC numbers, which is unique to IκB^−^ donor cell transplants and not seen in non-transplanted IκB^−^ mice, suggests that LT-HSC-autonomous NFκB activity limits their expansion or maintenance, and can be opposed by NFκB activity in the non-hematopoietic compartment.

**Figure 7.**
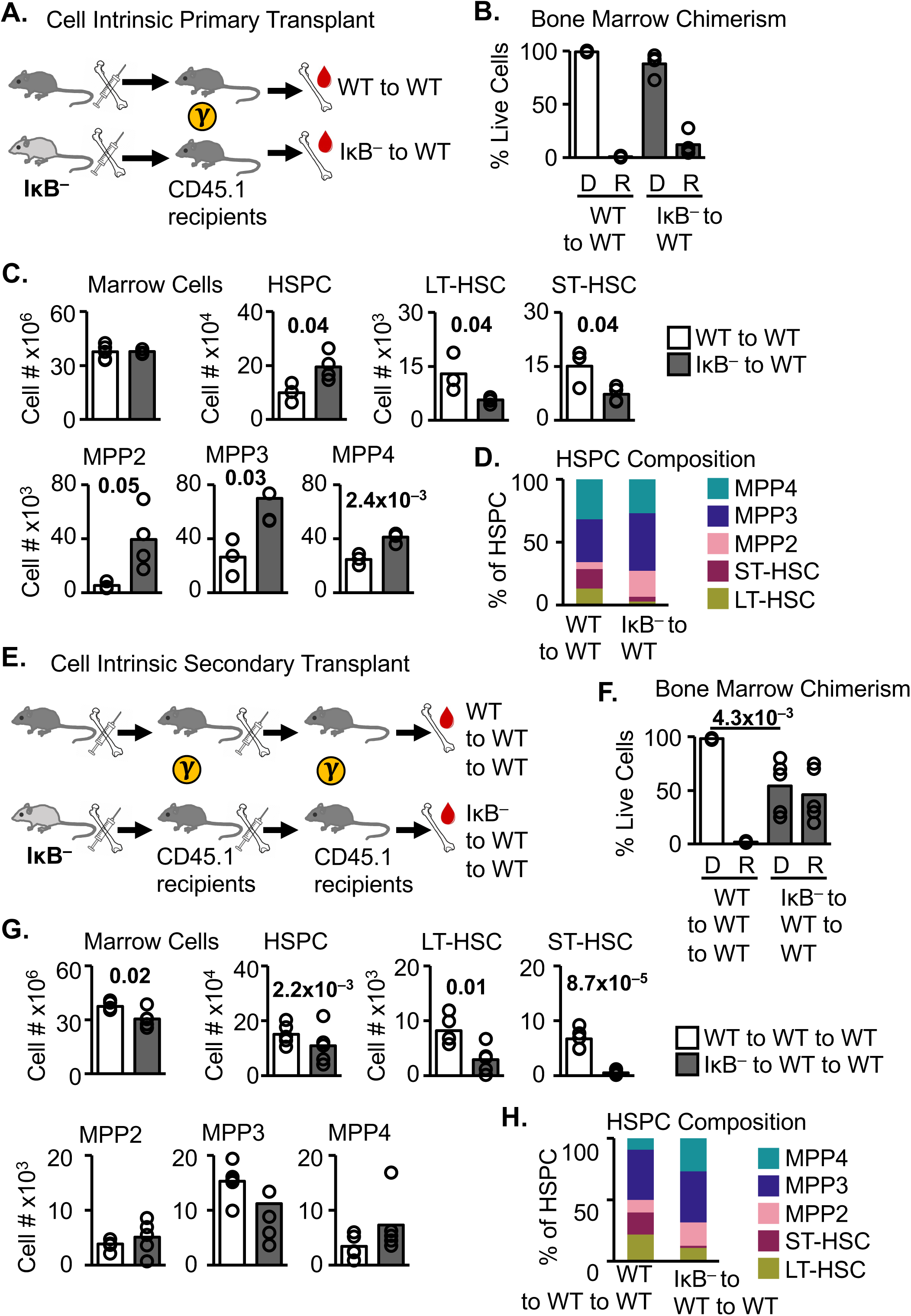
Experimentally elevated NFκB activity within HSCs impairs bone marrow reconstitution capacity. **A.** Primary cell-intrinsic transplant schematic. **B.** Bone marrow chimerism 15 weeks following primary transplant. **C.** Donor CD45.2 bone marrow, HSPC, and HSPC subset cell numbers. **D.** Donor CD45.2 bone marrow HSPC subsets (as in C), displayed as a percent of total donor HSPCs. **E.** Secondary cell-intrinsic transplant schematic. **F.** Bone marrow chimerism 16 weeks following secondary transplant; Donor (D), Recipient (R). **G.** Donor CD45.2 bone marrow, HSPC, and HSPC subset cell numbers. **H.** Donor CD45.2 bone marrow HSPC subsets (as in C), displayed as a percent of total donor HSPCs. A-D n=3-4; E-H n=5. Statistics calculated with unpaired students two-tailed t-test. See also Figure S7.

We further tested IκB^−^ HSC function by stressing their repopulation capacity using secondary bone marrow transplantation, which resulted in marked engraftment compromise of IκB^−^ donor cells compared to WT controls (peripheral blood chimerism 92% in WT donor vs 39% in IκB^−^ donor; bone marrow chimerism 98% in WT donor versus 54% in IκB^−^ donor) (**Figures S7C and 7E-F**). In addition, total bone marrow cellularity and total HSPC numbers were reduced (**Figure 7G**). Among HSPCs, LT-HSCs and ST-HSCs were both significantly reduced in number, and no lineage bias among MPP subsets was observed (**Figure 7G-H**). These findings demonstrate durable hematopoietic impairment 16 weeks following secondary transplantation. Further, the lack of HSPC expansion or MPP bias seen upon secondary transplantation of IκB^−^ donor marrow further supports that cell-autonomous NFκB activity is insufficient for myeloid bias. Together, these observations in IκB^−^ transplantation experiments indicate that elevated cell-intrinsic NFκB drives persistent functional HSC impairment with reduced long-term bone marrow reconstitution capacity, and is mechanistically separable from progenitor myeloid bias.

## DISCUSSION

By establishing the novel IκB^−^ model of elevated NFκB activity, we enabled the identification of distinct roles of NFκB in hematopoietic aging: in producing an inflammatory milieu that drives HSC epigenomic reprogramming and myeloid bias, and within HSCs in promoting their quiescence and limiting functional bone marrow reconstitution. At the same time, we found that HSC overabundance, another hallmark of hematopoietic aging, is not driven by NFκB. Importantly, these findings demonstrate that while myeloid bias, HSC dysfunction, and HSC accumulation co-occur in aging, they are mechanistically separable and are coordinated by different cellular compartments.

A key strength of the IκB^−^ model is its broad applicability to diverse states of inflammation. In the bone marrow, HSPCs, mature hematopoietic cells, and niche cells are all sources of inflammatory mediators in aging.^75,80^ Additionally, chronic inflammation may originate from systemic sources such as the microbiota, or in association with infection, cancer, autoimmune/inflammatory disease, and others.^81–83^ Despite variation in the quality, strength and cellular sources of inflammatory signaling between aging individuals, the phenotypes of myeloid bias, HSC dysfunction, and HSC accumulation are remarkably consistent. By targeting NFκB, a central regulatory node that integrates acute and chronic inflammatory signals, the IκB^−^ mouse is a robust model with relevance to a variety of inflammatory mediators and cellular sources, and to diverse individuals and underlying pathologies, unlike models of tissue-specific dysregulation of inflammatory mediators. IκB^−^ mice display myeloid bias and HSC dysfunction similar to aged mice. In contrast, this model demonstrates that systemically elevated NFκB activity is insufficient to drive the HSC accumulation seen in aging. Indeed, HSC pool size varies considerably between other models of systemic inflammation that display myeloid bias.^1^ These observations suggest that HSC accumulation may require additional perturbations such as DNA damage, metabolic derangements, or others.^84–87^ In future studies, models such as IκB^−^ may be useful to interrogate a wide range of chronic inflammatory states affecting HSCs beyond aging, such as myeloid neoplasia, bone marrow failure, iatrogenic immunomodulation, and more.^88–91^

NFκB activity exerts different effects in different cell types,^92–94^ and our findings clarify the functional pleiotropism of NFκB in HSCs versus their environments. We show here that NFκB activity in the bone marrow milieu drives myeloid bias, whereas HSC-intrinsic NFκB activity limits their functional bone marrow reconstitution capacity. Appreciating this diversity in NFκB-mediated functions sheds new light on the seemingly contradictory effects attributed to several cytokines and growth factors on hematopoiesis. TNF is elevated in the bone marrow microenvironment in both IκB^−^ and aged mice. Yet, TNF signaling (mediated through NFκB) can both restrict HSC proliferation/reconstitution and promote HSC expansion and myeloid bias.^95–97^ Our findings raise the possibility that the reduction in HSC reconstitution capacity following acute TNF may be mediated by HSC-intrinsic responses, while myeloid bias following chronic TNF exposure may be mediated through the milieu. IL-1 signaling is similarly appreciated to promote myeloid bias through a niche-mediated mechanism.^21,98^ Colony stimulating factors produced by the niche are NFκB-inducible,^23–25^ and an endothelial cell NFκB-CSF3 axis is recognized in emergency myelopoiesis.^99,100^ Concordantly, we found increased CSF3 in aged bone marrow, as well as increased CSF3R transcriptional responses in both aged HSCs and those transplanted into IκB^−^ recipients. However, while niche-derived acute CSF3 in emergency myelopoiesis is a potent inducer of myeloid differentiation and HSC mobilization, chronic CSF3 signaling is instead associated with HSC-autonomous quiescence and functional decline in a TLR/MyD88-dependent manner.^101,102^ TLR/MyD88 signaling is mediated in large part through NFκB,^22^ and thus our finding that NFκB mediates HSC quiescence and reconstitution impairment may reconcile these apparently opposing effects of CSF3 signaling in acute vs. chronic settings. As such, our contributions to understanding the pleiotropic roles of NFκB signaling in HSCs and their environments brings clarity to on-going debate about the complex roles of cytokines and growth factors in early hematopoiesis.

The results presented here also reconcile conflicting observations in the literature regarding whether myeloid bias is driven through cell-intrinsic or milieu-directed mechanisms. In aging, inflammation and myeloid bias co-occur, leading to initial assumptions that these phenotypes were co-driven in a cell-intrinsic manner.^28^ In support of this view, aged LT-HSCs have been shown to harbor cell autonomous transcriptional programs of myeloid bias and respond to inflammatory cues, impacting their differentiation.^11,75,103,104^ However, there has also been mounting evidence for niche-directed mechanisms of myeloid bias mediated by the sympathetic nervous system, vascular networks, mesenchymal lineage cells, and others.^28,59,105–111^ Our own recent work has also described milieu-directed myeloid bias requiring both differentiation bias and progenitor proliferation using mathematical models of early hematopoiesis (Singh et al., *Blood* accepted manuscript). Here, our studies demonstrate that inflammation in the bone marrow milieu is sufficient to drive phenotypic, transcriptomic and epigenomic myeloid bias of LT-HSCs, similar to those in aging. Thus, by identifying milieu-directed inflammation as a driver of HSC-intrinsic epigenomic and transcriptomic reprogramming, our findings reconcile the prior data in support of both HSC-extrinsic and HSC-autonomous mechanisms of myeloid bias.

HSC epigenomic programming can direct the cell fates of their progeny.^112^ Here, we identified premature or inappropriate epigenomic and transcriptomic myeloid signatures in HSCs underlying myeloid bias phenotypes, and which are induced by hematopoietic-extrinsic NFκB activity. A newly developed consistent cell labeling methodology enabled comparisons across omics modalities, datasets and species to generate these insights. In an inflamed niche, HSCs showed epigenomic reprogramming with increased C/EBP motif accessibility. This is consistent with results from an LPS-induced inflammation model;^113^ yet our findings further demonstrate that HSC epigenomic reprogramming is directed by an inflammatory milieu, rather than via HSC-autonomous responses. Lineage-primed MPPs are immediate HSC progeny; while they do not self-renew, they may transdifferentiate, with phenotypic megakaryocyte/erythroid-primed MPP2s and lymphoid-primed MPP4s contributing to myeloid output in inflammatory and regenerative contexts.^8,84^ Intriguingly, our study also identified potentiated C/EBP and PU.1 TF accessibility in MPP4s, raising the possibility that HSC epigenomic reprogramming may be heritable and contribute to apparent MPP transdifferentiation. This would be in-line with previous studies recognizing that C/EBP ectopic overexpression and high PU.1 favor myeloid cell fates over lymphoid.^8,45,49,114^ Here, we establish that increased C/EBP activity occurs in a natural setting of MPP differentiation bias, clarifying that the previous overexpression data is indeed physiologically relevant. Our findings from secondary hematopoietic-extrinsic transplantation experiments further show that phenotypic myeloid bias among multipotent progenitors is reversible upon return of HSCs to a non-NFκB-dysregulated microenvironment, despite the observed HSC epigenomic reprogramming that occurs during primary transplantation. This finding indicates that persistent inflammatory dysregulation is required to maintain myeloid bias phenotypes. Together, these omics data support HSCs as an important reservoir of immunological memory that is modifiable by milieu-derived signals, further pointing the bone marrow niche as a potential therapeutic target for modulating inflammatory dysregulation of hematopoiesis.^106,115^

In contrast to milieu-directed myeloid differentiation, we show here that HSC-intrinsic IκB^−^ promotes stem cell quiescence and diminished function similar to that seen in aging, chronic infection, and repeated inflammatory challenge.^1,6,116,117^. While paracrine mediators of HSC quiescence and maintenance are produced by local niches,^7,118^ our findings indicate an additional HSC-intrinsic mechanism of quiescence regulation. Previously, NFκB pathway activation *via* LTβR-agonism or IKK activating mutations provided conflicting evidence in regards to its impact on HSC quiescence.^119,120^ However, unlike the IκB^−^ model, these prior models modulate pathways other than NFκB, such as MAPK, autophagy and others.^29^ Here, we observed that NFκB activity and quiescence signatures are correlated in HSCs across the age-span, with both NFκB and quiescence signatures being potentiated in aging. These findings are compatible with the previously recognized steady-state dependence of HSCs on NFκB signaling,^121^ and with the sensitivity of aged HSCs to elevated NFκB signaling that limits their self-renewal.^122^ In other adult stem cells, chronic inflammation also limits stem cell exit from quiescence.^5^ These per-cell analyses provide insight into the heterogeneity within the stem cell pool, enabling new appreciation that transcriptional programs of myeloid bias are independent of HSC-intrinsic inflammatory NFκB-responses, and further support the framework of milieu-directed myeloid bias.

Phenotypic HSC heterogeneity has been recognized by cell surface expression of CD150, separating CD150^high^ myeloid-biased HSCs (myHSCs) from CD150^low^ lymphoid-biased HSCs (lyHSCs).^40,41^ MyHSCs accumulate with aging,^40,41^ an observation we reproduce here. However, we did not detect either an accumulation of myHSCs nor shift in myHSC vs. lyHSC phenotype in IκB^−^ mice. Recently, Semaphorin 4A has been identified as an important mediator of myHSC resistance to inflammatory stress in non-cell-autonomous manner.^79^ Thus, additional non-NFκB-driven mechanisms may control myHSC vs. lyHSC phenotype specification, in addition to the milieu-directed epigenomic and transcriptional myeloid bias detailed here. As myHSCs have been proposed as a target to improve suppressed lymphoid development and viral responses in aging,^123^ further work examining the regulation of HSC heterogeneity in steady-state, aging, and stress conditions will be important.

Our findings establish that inflammation-driven myeloid bias and HSC dysfunction are separable phenomena. HSC-intrinsic NFκB promotes HSC quiescence and limits stem cell function, while in contrast, an inflamed milieu directs HSC epigenomic reprogramming and myeloid bias. Taken together, these observations raise the possibility that distinct NFκB functions within HSCs and their cellular milieu may function to balance niche-derived emergency hematopoietic signals with HSC-driven quiescence and preservation. Our work indicates that HSCs and their environments should be considered independently to develop effective pharmacological strategies targeting NFκB and other signaling pathways to improve hematopoiesis in the setting of immune dysregulation.

## Resource Availability

Single cell RNA-sequencing and ATAC-sequencing data generated in this study are on GEO under accession numbers GSE261946 and GSE264569. Code and processed data objects will be made available following peer review.

## Acknowledgements

The authors thank Kylie Farrell, the UCLA JCCC, BSCRC, TCGB, IAC, and TPCL core facilities for technical assistance, and Quen Cheng for feedback on the manuscript. This research was funded by NIH R01CA264986 to DR and NIH R01AI127867, R01AI173214 and R01AI132731 to AH. JJC acknowledges the support of the California Institute for Regenerative Medicine UCLA Eli and Edythe Broad Center of Regenerative Medicine and Stem Cell Research Training Program, and AS acknowledges training grant support from NIH T32GM008185 and T32GM008042.

## Authorship Contributions

Conceptualization, J.J.C. and A.H.; Methodology, J.J.C, A.S, Y.S.L, Y.L., and J.K.K.; Investigation, J.J.C, Y.S.L, Y.L, T.T. and J.K.K.; Formal Analysis, J.J.C, A.S., N.P. and D.M.; Resources, D.S.R and A.H.; Writing, J.J.C. and A.H.; Supervision, D.S.R and A.H.; Funding Acquisition, D.S.R and A.H.

The current affiliation for Yi Liu is DeepKinase Biotechnologies, Ltd, Beijing, China. The current affiliation for Noa Popko is University of California, San Diego, Department of Bioinformatics and Systems Biology.

## Supplemental Information Index

Supplemental Figures S1-7, related to Figures 1-7

Table S1, Gene lists used to calculate PCA and single-cell enrichment scores. Excel file containing additional data too large to fit in a PDF, related to Figures 4-6.

Tables S2, Gene ontology results from Quiescence^high^ versus Myeloid^high^ downregulated DEG in aged LT-HSCs. Excel file containing additional data too large to fit in a PDF, related to Figure 6.

Table S3, Gene ontology results from Quiescence^high^ versus Myeloid^high^ upregulated DEG in aged LT-HSCs. Excel file containing additional data too large to fit in a PDF, related to Figure 6

## MATERIALS AND METHODS

### Mice

Young (2-3 months) and aged (18-22 months) male and female C57Bl/6 and B6.SJL-Ptprc^a^Pepc^b^/BoyJ (CD45.1) mice were obtained from The Jackson Laboratory and our own breeding colony. Previously described *Nfkbia*^+/-^ mice^34^ were bred with *Nfkbib*^-/-^ and *Nfkbie*^-/-^ strains^38,39^ to generate “IκB^−^” *Nfkbia^+/-^Nfkbib^-/-^Nfkbie^-/-^*. mVenus-RelA mice were previously described.^35^ Animals were maintained and experiments conducted in accordance with protocols approved by the UCLA Institutional Animal Care and Use Committee.

### Flow cytometry and cell sorting

Peripheral blood and bone marrow cells collected, processed, stained with fluorochrome-conjugated antibodies, and HSPC cell populations identified as previously described.^8,124–126^ For cell sorting, lineage-positive cells were pre-depleted using Invitrogen MagniSort™ Mouse Hematopoietic Lineage Depletion Kit (Invitrogen, Cat#8804-6829-74) per the manufacturer protocol, with lineage-biotin antibody cocktail at 100μl/10 million cells, and SA-magnetic beads at 20μl/10 million cells. Flow cytometry performed with a LSRFortessa X-20 (BD Biosciences). Flow sorting performed with FACS Aria™III Cell Sorters (BD Biosciences). Cells counted via CytoFlex (Beckman Coulter). Analysis performed using FlowJo v10.8.1 (TreeStar).

### Cell quantification from flow cytometry

To calculate cell number, the percentage of the gated population was multiplied by the absolute cell counts.^124^ To calculate HSPC composition, the percentage of the gated population was calculated by dividing the number of cells within a subset (e.g. LT-HSC) by the total number of HSPCs (Lin^−^/Sca1^+^cKit^hi^).

### Histology and microscopy

Mouse sterna were formaldehyde-fixed and paraffin-embedded per standard histological protocols, then sectioned and stained with H&E. Analysis performed by a board-certified Hematopathologist (J.J.C.), and photographed using a BX43 microscope, DP27 camera and cellSens Standard 3.2 (all Olympus).

### Luminex

Bone marrow from bilateral femurs was collected via centrifugation^126^ in 200μl PBS; cells were resuspended in the supernatant and re-pelleted at 5000g for 3mins at 4C. Supernatants were immediately collected and stored at −80C. At the time of assay, samples were thawed, spun at 13,000xg for 10 minutes at 4C, and low-input format mouse 32-plex Luminex (EMD Millipore) was performed per manufacturer’s protocols.

### Bone marrow transplantation

Bone marrow transplantation was performed as previously described.^125^ Briefly, recipient mice were lethally irradiated with a Cesium irradiator (1100 rads); 24 hours later, 3-5 million total donor bone marrow cells were injected retro-orbitally. Animals were maintained in sterile cages with sterilized food and autoclaved water for two weeks following irradiation. Thereafter, animals were maintained in autoclaved cages. Peripheral blood chimerism was evaluated at the specified time points by retro-orbital bleed.

### ATAC-seq

#### Cell isolation and library preparation

CD45.1 donor LT-HSC, MPP2, MPP3 and MPP4 populations were sorted from WT or IκB^−^ recipients 13-15 weeks after transplantation, with 650 −10,000 cells/population. Nuclei isolation, transposase reactions, and library preparation were performed as previously described,^127^ separately for n=3 pairs of chimeric mice.

#### Sequencing and pre-processing

Libraries were multiplexed and single-end sequenced with a length of 75bp on the NextSeq500 High Output sequencing system. Read trimming, alignment, filtering, duplicate removal, peak calling and browser track preparation were performed as previously described.^127^ Consensus peaks (13910) were identified between biological replicates using ChIP-R v1.1.0.^128^

#### Analysis

Differential accessibility analysis was performed in R v4.3.2 with EdgeR v4.0.9 using TMM normalization, glmQLFit and glmTreat arguments.^129^ TF accessibility analysis and peak annotations were performed using Homer v4.10.3.^130^ ChIP datasets C/EBPα^51^ (GSM1347229), C/EBPβ^52^ (GSM2974656) and C/EBPδ^53^ (GSM2663838). ATAC-seq and ChIP-seq peak overlaps identified using GenomicsRanges v1.54.1.^131^ Consensus peaks visualized using Integrative Genomics Viewer v2.9.4.^132^ For comparison to scRNA-seq, peaks were assigned to the nearest gene.

### scRNA-seq

#### Cell isolation and library preparation

CD45.1 donor HSPCs (Lin^−^/Sca1^+^cKit^hi^) were sorted from WT or IκB– recipients 15 weeks after transplantation. ∼30,000 HSPCs were incubated with TotalSeq-B barcode antibodies Hashtag#1 (WT recipient) and #2 (IκB^−^ recipient; Biolegend, Cat#155831,155833 respectively) per manufacturer’s protocol, then pooled for sequencing. Single cell transcriptome and barcode libraries were generated using Chromium Next GEM Single Cell 3 Reagent Kits v3.1 (10X Genomics).

#### Sequencing and pre-processing

Libraries were paired-end sequenced with a length of 50bp on the NovaSeqSP system. Read counts were quantified using Cellranger v7.0.^133^ Standard quality-control criteria based on minimum UMI, percent mitochondrial counts, and potential doublets were applied. Principal component analysis (PCA) was calculated from 115 hematopoietic genes (Table S1).

#### HSPC labeling model

Microarray reference data from flow-sorted LT-HSC, ST-HSC, MPP2, MPP3 and MPP4^8^ was filtered by reference flow-sorted scRNA-seq data^54^ identifying13522 reference probes mapping to 11644 reference scRNA-seq genes with nonzero counts in >1% of cells. A logistic regression model using glmnet v4.1-7^134^ was built by subsampling 1% of filtered reference probes for 1000 iterations imposing an L1 lasso penalty, and limited to non-negative values to identify positive marker genes. The mean of the iterations was calculated yielding a final labeling coefficient matrix of 8511 probes mapping to 7563 genes. To apply the labeling model, final coefficient matrix was multiplied by the scRNA-seq genes by cells matrix, yielding scores for all 5 cell types across every cell in the dataset. Cell-type probabilities were calculated by dividing each cell type score by the sum of all 5 scores within a single cell; the predicted cell identity was then set to the highest probability. For human datasets, ribosomal genes were excluded.^135^

#### Analysis

Principal components were calculated using 115 genes with established roles in hematopoiesis. DEG analysis (Figure 4) performed using Seurat v4.4.0.^136^ Single cell gene set enrichment scores calculated as normalized mean rank for each gene set and z-scored by dataset.^55^ TF activity estimated using decoupleR v2.8.0.^64^ Aged DEG pseudobulk analysis performed by summing single cell counts in EdgeR v3.3.6 with glmFit and glmLRT, and experiments as a covariate.^129^ Fgsea v1.2.0 used to generate GO ranked by logFC and collapsePathways, with terms from msigdb v7.5.1.^137,138^ Human to mouse genes mapped with bioMart v2.5.0. All gene lists available in Table S1.

### Data and resource sharing

scRNA-seq and ATAC-seq data generated in this study available at GEO under accession numbers GSE261946 and GSE264569. Cell labeling code and processed data objects available at https://doi.org/10.5281/zenodo.10828398. Further information and requests for resources and reagents will be fulfilled by the corresponding author.

### Quantification and statistical analysis

Statistical testing methodologies are specified for each analysis in the relevant figure legend. T-tests were performed in Excel (Microsoft). Wilcoxon Rank Sum tests, KS tests of the distribution, and Spearman correlations were performed in R v4.3.2. ATAC-seq TF enrichment analyses employed ZOOPS scoring in Homer v4.10.3. BH adjusted p-values for Figure 4 DEG performed using Seurat v4.4.0. BH adjusted p-values for Figure 6 generated in EdgeR v3.3.6 with glmFit and glmLRT, and experiments as a covariate. Where unreported, p-value is > 0.05.

## RESOURCE TABLE

| REAGENT or RESOURCE | SOURCE | IDENTIFIER |
| --- | --- | --- |
| **Antibodies** |  |  |
| CD135 (Flt3) Monoclonal Antibody (A2F10), PerCP-eFluor 710, eBioscience™ | Invitrogen | Cat# 46-1351-82 ;<br>RRID: AB_10733393 |
| CD3 Monoclonal Antibody (17A2), PerCP-eFluor 710, eBioscience™ | Invitrogen | Cat# 46-0032-82 ;<br>RRID: AB_1834427 |
| Purified anti-mouse CD16/32 Antibody | BioLegend | Cat# 101302; RRID:<br>AB_312801 |
| PE/Dazzle™ 594 anti-mouse/human CD11b Antibody | BioLegend | Cat# 101256; RRID:<br>AB_2563648 |
| APC anti-mouse/human CD45R/B220 Antibody | BioLegend | Cat# 103212; RRID:<br>AB_312997 |
| PE/Cy7 anti-mouse Ly-6G/Ly-6C (Gr-1) Antibody | BioLegend | Cat# 108416; RRID: AB_313381 |
| Brilliant Violet 711™ anti-mouse CD45.1 Antibody | BioLegend | Cat# 110739; RRID: AB_2562605 |
| FITC anti-mouse CD45.1 Antibody | BioLegend | Cat# 110706; RRID: AB_313495 |
| APC/Fire™ 750 anti-mouse CD45.1 Antibody | BioLegend | Cat# 110752; RRID: AB_2629806 |
| Brilliant Violet 711™ anti-mouse CD45.2 Antibody | BioLegend | Cat# 109847; RRID: AB_2616859 |
| FITC anti-mouse CD45.2 Antibody | BioLegend | Cat# 109806; RRID: AB_313443 |
| PE anti-mouse TER-119/Erythroid Cells Antibody | BioLegend | Cat# 116208 ; RRID: AB_313709 |
| PE anti-mouse CD117 (c-Kit) Antibody | BioLegend | Cat# 105808 ; RRID: AB_313217 |
| PE/Cy7 anti-mouse Ly-6A/E (Sca-1) Antibody | BioLegend | Cat# 108114 ; RRID: AB_493596 |
| APC/Cyanine7 anti-mouse CD48 Antibody | BioLegend | Cat# 103432; RRID: AB_2561463 |
| APC anti-mouse CD150 (SLAM) Antibody | BioLegend | Cat# 115910; RRID: AB_493460 |
| Brilliant Violet 711™ anti-mouse CD117 (c-Kit) Antibody | BioLegend | Cat# 105835; RRID: AB_2565956 |
| Brilliant Violet 510™ Streptavidin | BioLegend | Cat# 405234; N/A |
| Brilliant Violet 711™ Streptavidin | BioLegend | Cat# 405241; N/A |
| TotalSeq-B barcode antibodies Hashtag#1 | BioLegend | Cat# 155831; AB_2814067 |
| TotalSeq-B barcode antibodies Hashtag#2 | BioLegend | Cat# 155833; AB_2814068 |
| Chemicals, peptides, and recombinant proteins |  |  |
| DAPI (4',6-Diamidino-2-Phenylindole, Dilactate) | Invitrogen | Cat# D3571; RRID: AB_2307445 |
| Critical commercial assays |  |  |
| Luminex mouse 32-plex assay | EMD Millipore | Cat# MCYTMAG-70K-PX32; N/A |
| MagniSort™ Mouse Hematopoietic Lineage Depletion Kit | Invitrogen | Cat# 8804-6829-74; AB_2575269 |
| Chromium Next GEM Single Cell 3' GEM, Library & Gel Bead Kit v3.1 | 10X Genomics | Cat# PN-1000121; N/A |
| Illumina Tagment DNA Enzyme and Buffer Small Kit | Illumina | Cat# 20034197; N/A |
| Nextera DNA Library Preparation Kit (24 samples) | Illumina | Cat# FC-121-1030; N/A |
| MinElute PCR Purification Kit (50) | Qiagen | Cat# 28004; N/A |
| Library Quant Kit (Illumina, Universal) | KAPA Biosystems | Cat# KK4824; N/A |
| Deposited data |  |  |
| Murine HSPC subset bulk ATAC-sequencing data | This manuscript | GSE264569 |
| Murine HSPC single-cell RNA sequencing data | This manuscript | GSE261946 |
| Murine ChIP for C/EBPα | Jakobsen et al.<sup>51</sup> | GSM1347229 |
| Murine ChIP for C/EBPβ | Link et al.<sup>52</sup> | GSM2974656 |
| Murine ChIP for C/EBPδ | Comoglio et al.<sup>53</sup> | GSM2663838 |
| Murine HSPC subset sorted bulk microarray data | Pietras et al.<sup>8</sup> | GSE68529 |
| Murine HSPC subset sorted single-cell RNA sequencing data | Rodriguez-Fraticelli et al.<sup>54</sup> | GSE90742 |
| Murine HSPC single-cell RNA sequencing data | Mitchell et al.<sup>59</sup> | GSE169162 |
| Murine HSPC single-cell RNA sequencing data | Hérault et al.<sup>60</sup> | GSE147729 |
| Human CD34-positive single-cell RNA sequencing data | Ainciburu et al.<sup>61</sup> | GSE180298 |
| Human CD34-positive subset sorted single-cell RNA sequencing data | Zhang et al.<sup>62</sup> | GSE137864 |
| *Mus musculus* Genome assembly GRCm38/mm10 | NCBI, UCSC | https://www.ncbi.nlm.nih.gov/datasets/genome/GCA_000001635.2/<br>https://genome.ucsc.edu/cgi-bin/hgGateway?db=mm10 |
| Experimental models: Organisms/strains |  |  |
| Mouse: C57BL/6J | The Jackson Laboratory | JAX: 000664 ; RRID: IMSR_JAX:000664 |
| Mouse: B6.SJL-*Ptprc*<sup>a</sup> *Pepc*<sup>b</sup>/BoyJ | The Jackson Laboratory | JAX: 002014 ; RRID: IMSR_JAX:002014 |
| Mouse: RelA-mVenus (C57BL/6J background) | Adejala et al.<sup>35</sup> | N/A |
| Mouse: *Nfkbib*<sup>-/-</sup> *Nfkbie*<sup>-/-</sup> *Nfkbia*<sup>+/-</sup> (C57BL/6J background) | This manuscript | N/A |
| Software and algorithms |  |  |
| FlowJo | TreeStar | v10.8.1; N/A |
| R | https://www.r-project.org | v4.3.2; N/A |
| ChIP-R | Newell et al.<sup>128</sup> | v1.1.0; N/A |
| EdgeR | Robinson et al.<sup>129</sup> | v3.3.6 (scRNA-seq), v4.0.9 (ATAC-seq); N/A |
| HOMER Motif Analysis | Heinz et al.<sup>130</sup> | v4.10.3; N/A |
| GenomicsRanges | Lawrence et al.<sup>131</sup> | v1.54.1; N/A |
| Integrative Genomics Viewer | Robinson et al.<sup>132</sup> | v2.9.4; N/A |
| Cellranger | Zheng et al.<sup>133</sup> | V7.0; N/A |
| glmnet | Friedman et al.<sup>134</sup> | v4.1-7; N/A |
| Seurat | Butler et al.<sup>136</sup> | v4.4.0; N/A |
| decoupleR | Badia-i-Mompel et al.<sup>64</sup> | v2.8.0; N/A |
| Fgsea | Korotkevich et al.<sup>137</sup> | v1.2.0; N/A |
| msigdb | Castanza et al.<sup>138</sup> | v7.5.1; N/A |
| bioMart | Durinck et al.<sup>139</sup> | v2.5.0; N/A |
| Code, including HSPC labeling and scRNA-sequencing analysis | This manuscript | https://doi.org/10.5281/zenodo.10828398 |

## CONTACT FOR REAGENT AND RESOURCE SHARING

Requests for further information, resources and reagents will be fulfilled by the corresponding author.

### Abbreviations

ATAC-seq: Assay for transposase-accessible chromatin with sequencing
BH: Benjamini-Hochberg
CD: cluster of differentiation
CSF3: colony stimulating factor 3; aka G-CSF
CSF3R: colony stimulating factor 3 receptor; aka G-CSFR
C/EBP: CCAAT/enhancer family of basic-leucine zipper transcription factors
ChIP-seq: chromatin immunoprecipitation with sequencing
CPM: counts per million
DAMP: damage associated molecular pattern
DAR: differentially accessible chromatin region
DEG: differential gene expression
GO: gene ontology
H&E: hematoxylin and eosin
HSC: hematopoietic stem cell
HSPC: hematopoietic stem and progenitor cell
IκB: inhibitor of NFκB
IKK: IκB kinase
IL: interleukin
NFκB: nuclear factor kappa B
LTβR: lymphotoxin beta receptor
MPP: multipotent progenitor
MAP: mitogen activated protein
My: Myeloid
NIK: NFκB-inducing kinase
PAMP: pathogen associated molecular pattern
PI3: Phosphoinositide 3
Ly: Lymphoid
scRNA-seq: single cell ribonucleic acid sequencing
TF: transcription factor
TGFβ: tissue growth factor beta
TNF: tumor necrosis factor
WT: wild type

