## Supplemental Figures for "Distinct causes of three phenotypic hallmarks of hematopoietic aging"

**Supplemental Figure 1**

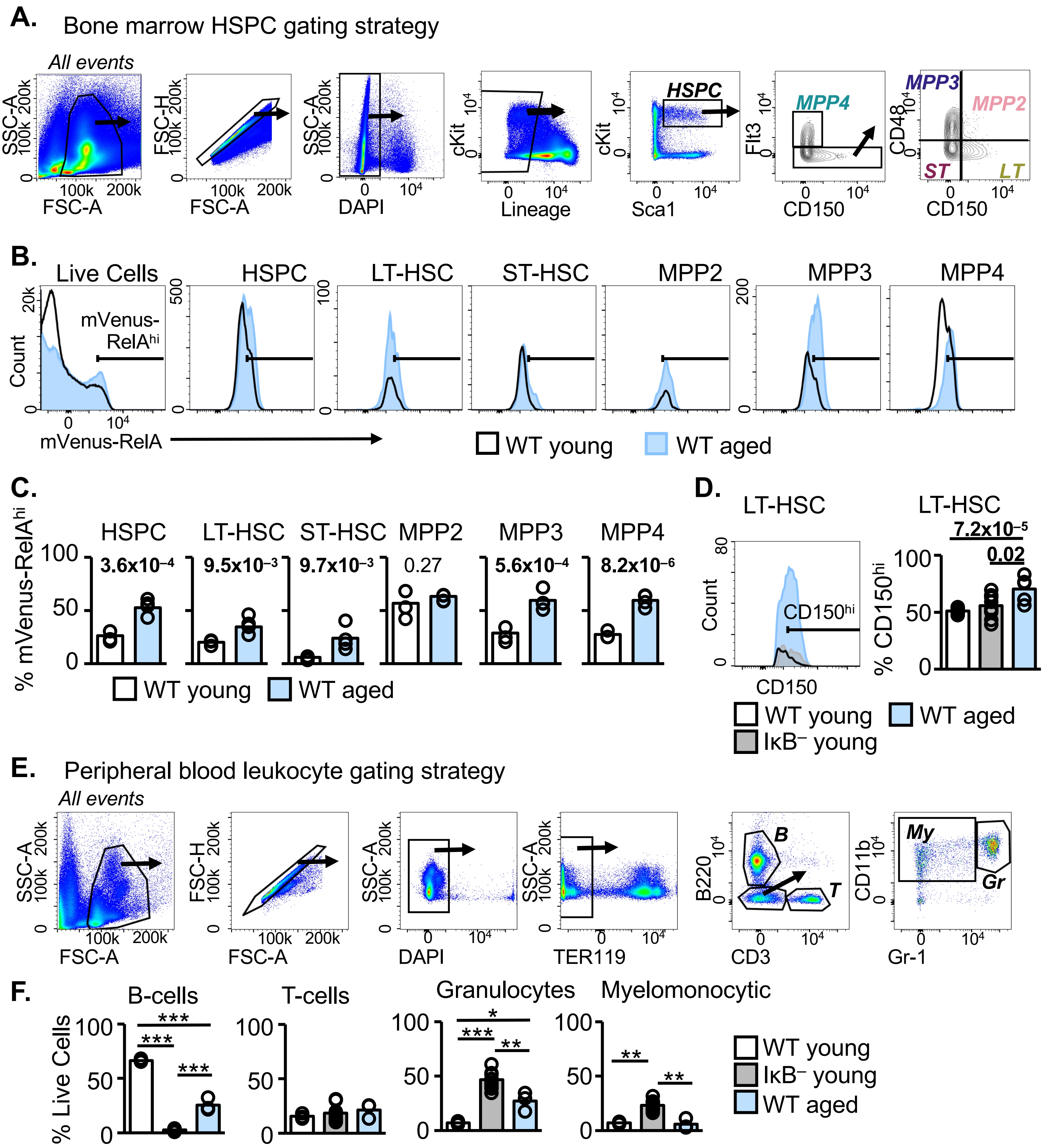


**Supplemental Figure 1. Flow cytometry analysis of murine bone marrow and peripheral blood. A.** Flow cytometry gating strategy for bone marrow HSPC and subsets. **B.** mVenus-RelA fluorescence by flow cytometry in the indicated HSPC subpopulations; gates indicate RelA$^{hi}$ threshold; representative of n=4-5. **C.** Quantification of RelA$^{hi}$ cells as gated in B, comparing young and aged for each subset. **D.** Representative histogram (left) and quantification (right) of CD150$^{hi}$ LT-HSCs in the indicated ages and genotypes; n=5-12. **E.** Flow cytometry gating strategy for peripheral blood leukocyte populations. **F.** Peripheral blood leukocyte composition by flow cytometry; n=3-11. Statistics calculated with unpaired students two-tailed t-test *p<0.05, **p<0.01, ***p<0.001. Where unreported, pvalue is > 0.05.

**Supplemental Figure 2**

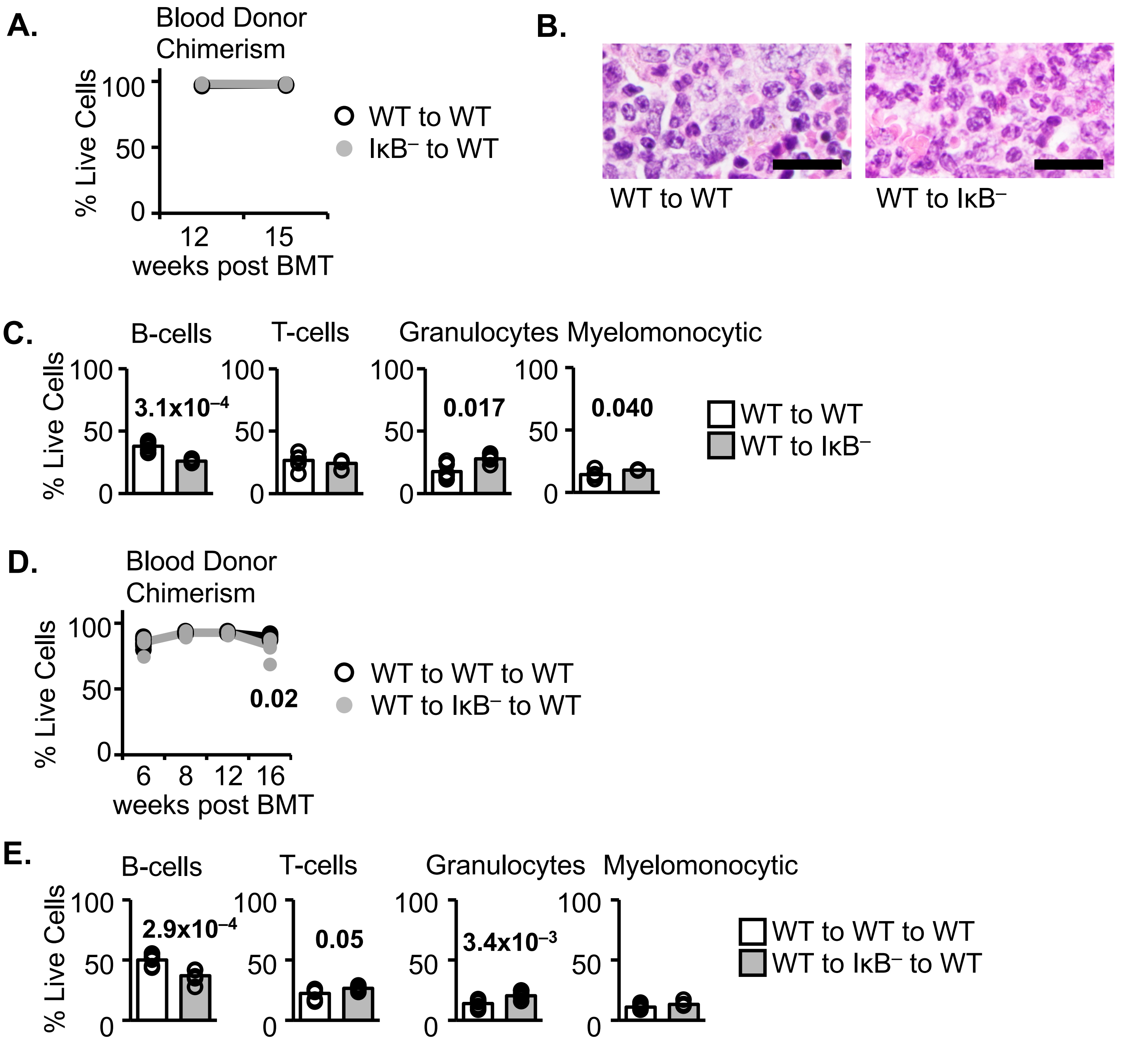


**Supplemental Figure 2. Cell-extrinsic bone marrow transplant peripheral blood analyses**.
**A.** Primary cell-extrinsic peripheral blood donor chimerism at the indicated weeks following primary transplant. **B.** Mature granulocytosis in sternum bone marrow histology 15 weeks following primary transplant, H&E; scale bars are 20μm. **C.** Donor CD45.1 peripheral blood leukocyte composition at 15 weeks following primary transplant. **D.** Secondary cell-extrinsic peripheral blood donor chimerism at the indicated weeks following secondary transplant. **E.** Donor CD45.1 peripheral blood leukocyte composition at 16 weeks following secondary transplant. A-C n=4-6; D-E n=7. Statistics calculated with unpaired students two-tailed t-test. Where unreported, pvalue is > 0.05.

**Supplemental Figure 3**

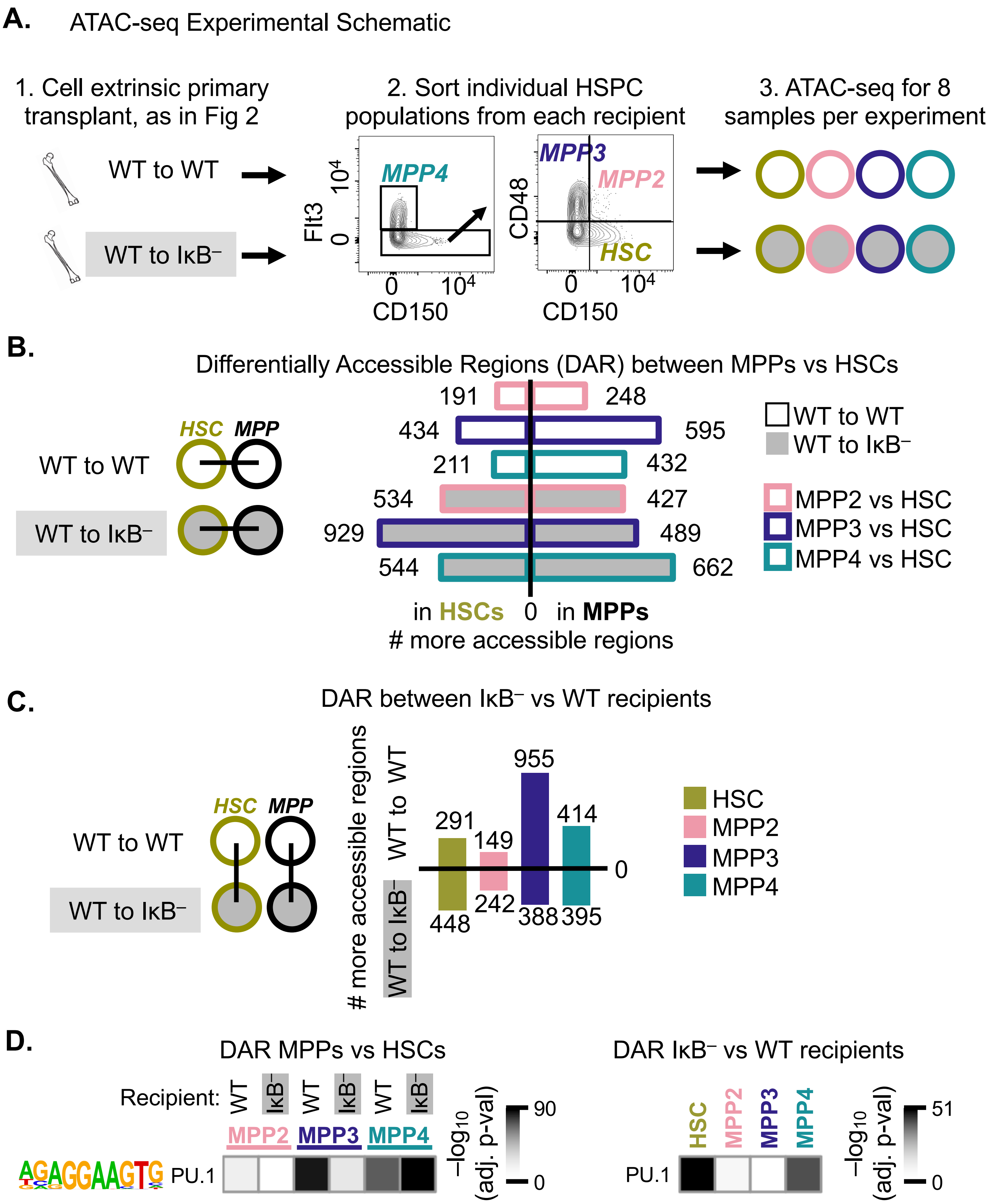


**Supplemental Figure 3. ATAC-seq experimental design and differential accessibility comparisons.** **A.** Experimental schematic. **B.** Number of differentially accessible chromatin regions (DAR) that are more accessible in the indicated populations, out of 13910 total regions; corresponds to analysis in Fig 3B. **C.** Number of DAR that are more accessible in the indicated populations, out of 13910 total regions; corresponds to analysis in Fig 3C. **D.** PU.1 TF motif enrichment.

**Supplemental Figure 4**

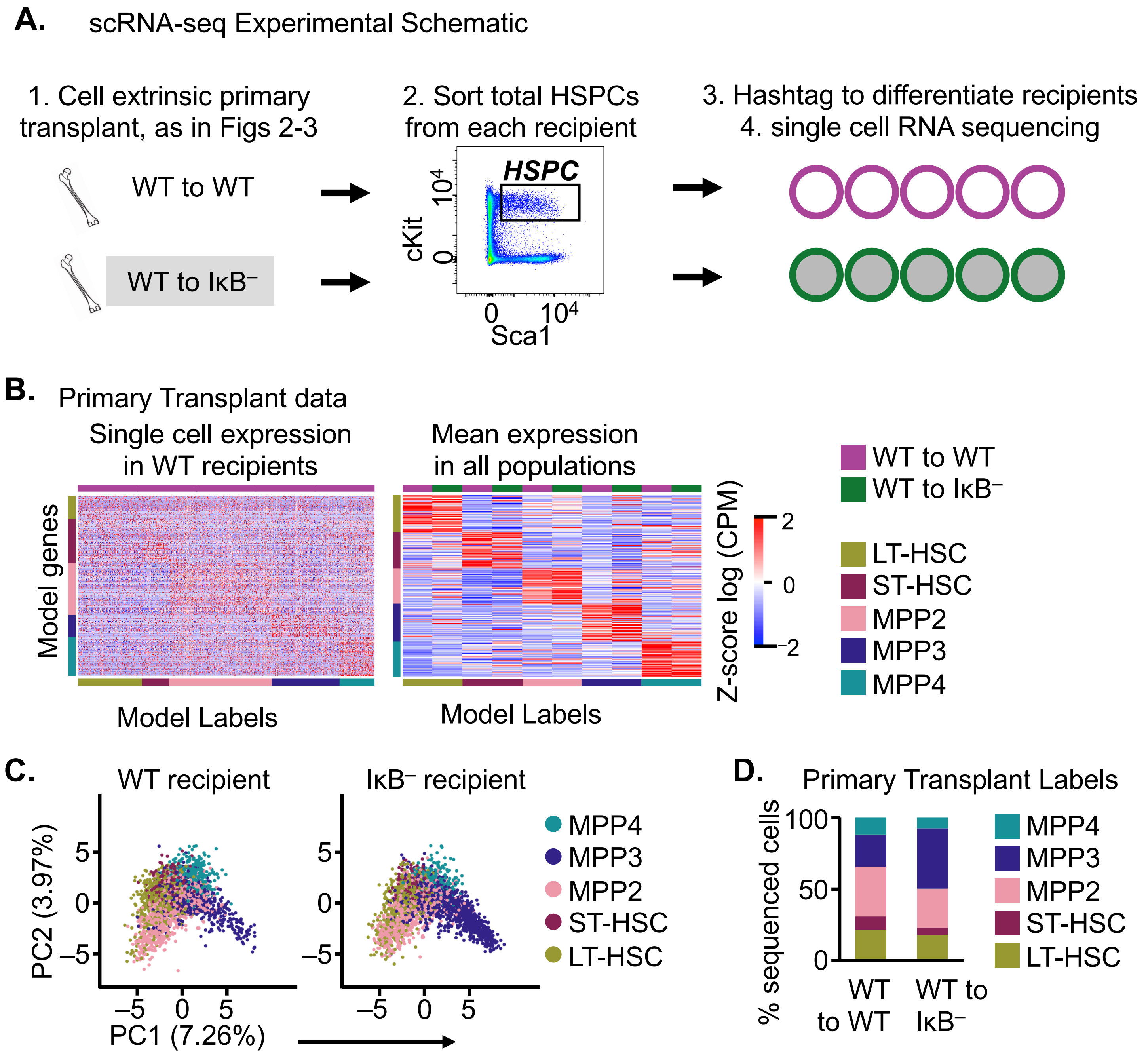


**Supplemental Figure 4. scRNA-seq experimental design and cell labeling. A.** Experimental schematic. **B.** Expression of the top 10% cell type-specific model genes in single cells with non-zero expression for >1% of the displayed genes (left) and for the top 10% cell type-specific model genes as a mean of each population (right). **C.** Data colored by cell identity assignment, split by recipient condition. **D.** Proportion of sequenced HSPC in each subset.

**Supplemental Figure 5**

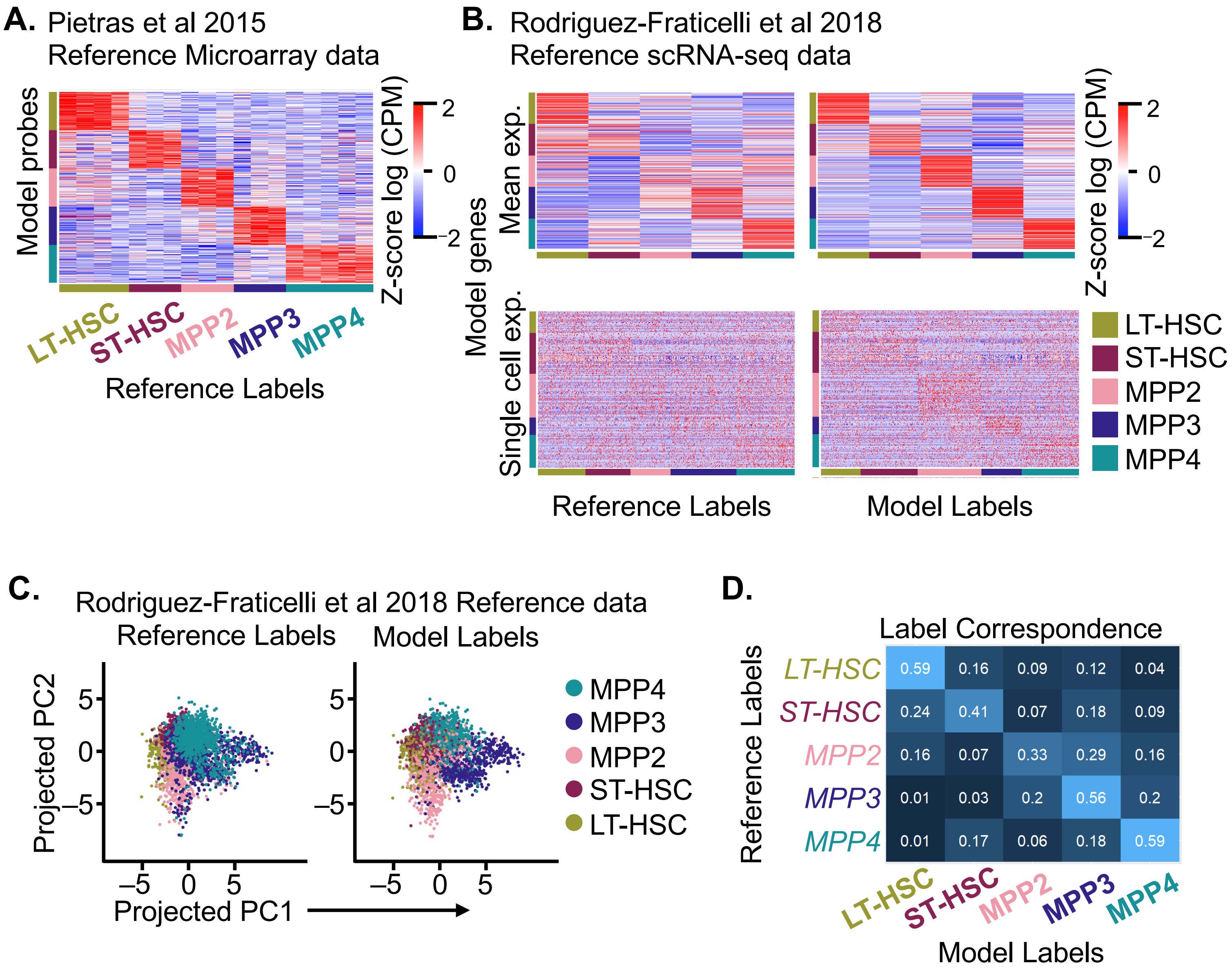


**Supplemental Figure 5. Data-driven HSPC labeling model. A.** Microarray expression of label model probes in all microarray samples. **B.** scRNA-sequencing expression of the top 10% cell type-specific model genes as a mean of the classified population (top) and in single cells with non-zero expression for >1% of the displayed genes (bottom), when cells are classified according to the reference labels (left) or the model assigned labels (right). **C.** scRNA-seq reference data projected onto PCA space from the transplant dataset, colored per the reference (left) or model labels (right). **D.** Correspondence between reference and model labels for each population.

**Supplemental Figure 6**

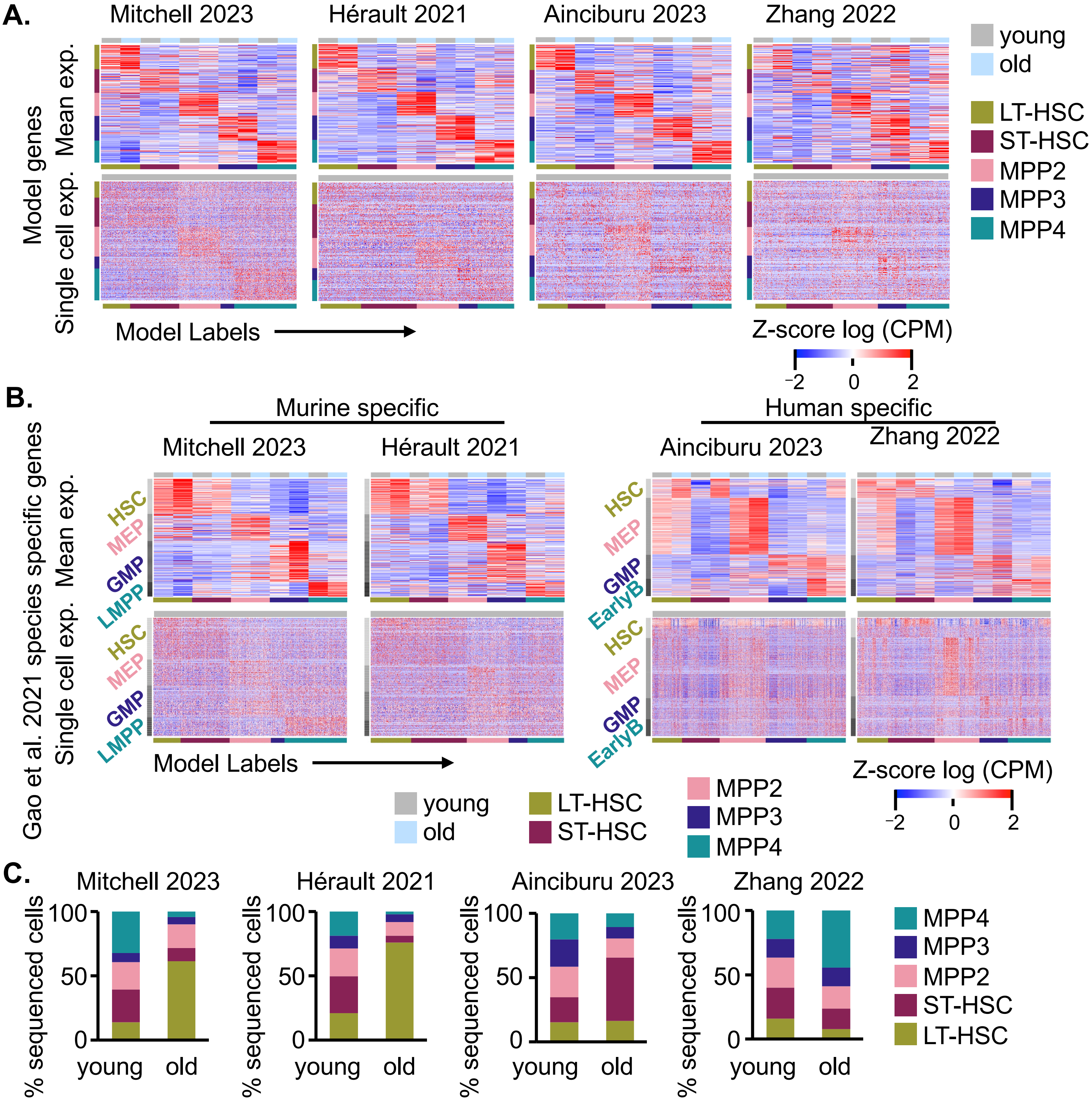


**Supplemental Figure 6. Model labeling of aged datasets. A.** Expression of the top 10% cell type-specific model genes as a mean of each population (top) and in single cells with non-zero expression for >1% of the displayed genes (bottom). **B.** Expression of species-specific genes identified by Gao and colleagues for the designated cell types, as a mean of each population (top) and in single cells (bottom). **C.** Proportion of sequenced HSPC in each subset.

**Supplemental Figure 7**

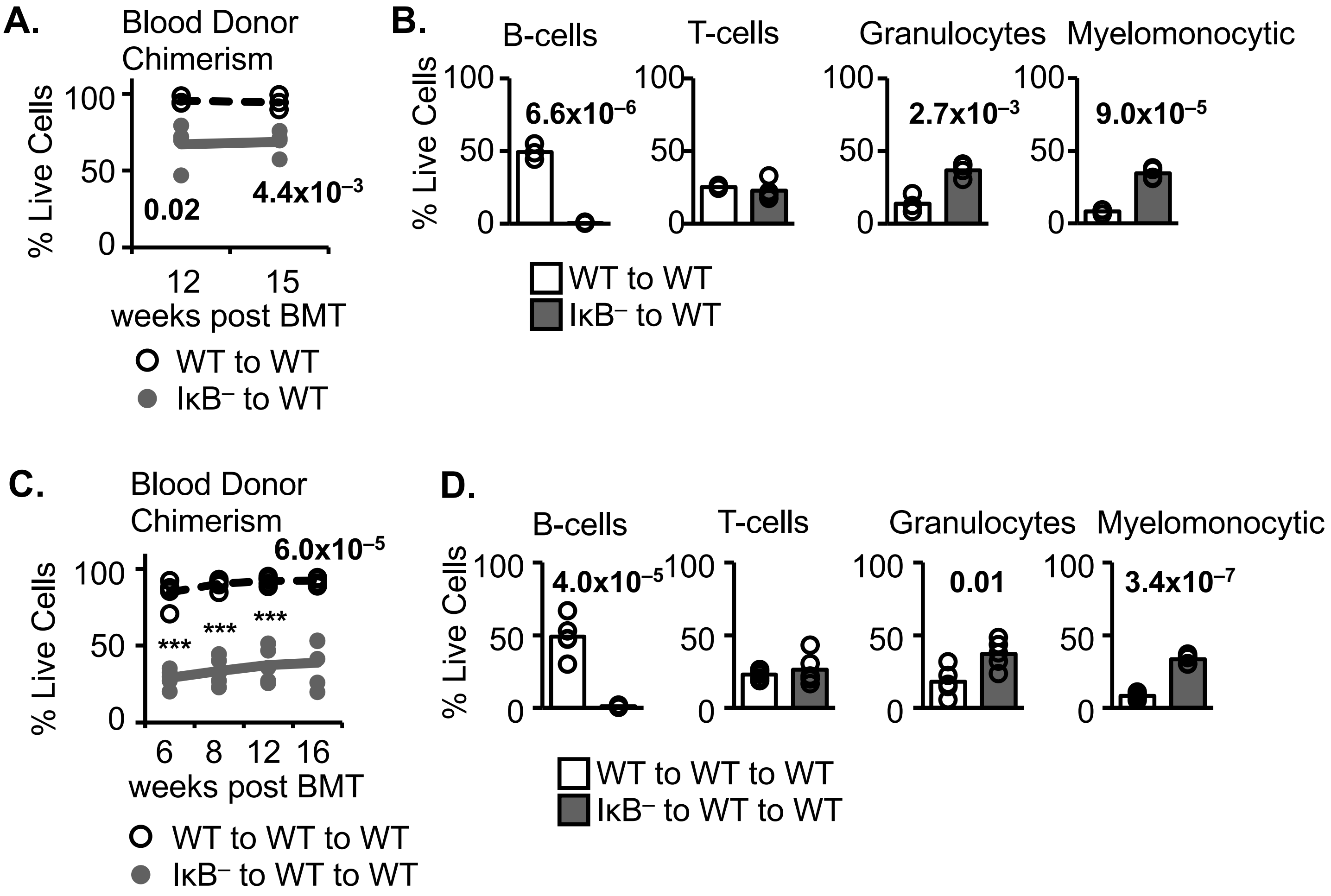


**Supplemental Figure 7. Cell-intrinsic bone marrow transplant peripheral blood analyses. A.** Primary cell-intrinsic peripheral blood donor chimerism at the indicated weeks following primary transplant. **B.** Donor CD45.2 peripheral blood leukocyte composition at 15 weeks following primary transplant. **C.** Secondary cell-intrinsic peripheral blood donor chimerism at the indicated weeks following secondary transplant. **D.** Donor CD45.2 peripheral blood leukocyte composition at 16 weeks following secondary transplant. A-B n=3-4; C-D n=5. Statistics calculated with unpaired students two-tailed t-test ***$p<1x10^{-6}$. Where unreported, pvalue is > 0.05.
